# Integrating convolution and self-attention improves language model of human genome for interpreting non-coding regions at base-resolution

**DOI:** 10.1101/2021.09.06.459087

**Authors:** Meng Yang, Haiping Huang, Lichao Huang, Nan Zhang, Jihong Wu, Huanming Yang, Feng Mu

## Abstract

Interpretation of non-coding genome remains an unsolved challenge in human genetics due to impracticality of exhaustively annotate biochemically active elements in all conditions. Deep learning based computational approaches emerge recently to help interpretating non-coding regions. Here we present LOGO (<u>L</u>anguage <u>o</u>f <u>G</u>en<u>o</u>me), a self-attention based contextualized pre-trained language model containing only 2 self-attention layers with 1 million parameters as a substantially light architecture that applies self-supervision techniques to learn bidirectional representations of unlabeled human reference genome. LOGO is then fine-tuned for sequence labelling task, and further extended to variant prioritization task via a special input encoding scheme of alternative alleles followed by adding a convolutional module. Experiments show that LOGO achieves 15% absolute improvement for promoter identification and up to 4.5% absolute improvement for enhancer-promoter interaction prediction. LOGO exhibits state-of-the-art multi-task predictive power on thousands of chromatin features with only 3% parameterization benchmarking against fully supervised model, DeepSEA and 1% parameterization against a recent BERT-based language model for human genome. For allelic-effect prediction, locality introduced by one dimensional convolution shows improved sensitivity and specificity for prioritizing non-coding variants associated with human diseases. In addition, we apply LOGO to interpret type 2 diabetes (T2D) GWAS signals and infer underlying regulatory mechanisms. We make a conceptual analogy between natural language and human genome and demonstrate LOGO is an accurate, fast, scalable, and robust framework to interpret non-coding regions for global sequence labeling as well as for variant prioritization at base-resolution.

---

In 2003, the Human Genome Project (HGP) has successfully digitalized the ‘book of life’. It is convinced that biological structure and function are intrinsically encoded in the primary genome sequence. The non-coding regions, accounting for over 98% of the whole genome, implement significant yet largely unknown regulatory functions. Recent large consortia projects, including the ENCyclopedia of DNA Elements (ENCODE)^1,2^, Roadmap Epigenomics^3^, and the Genomics of Gene Regulation (GGR), have produced large amount of experimental mapping readouts to help annotate non-coding genome in specific tissues or cell-lines. On the other hand, Genome-wide association studies (GWAS) have discovered that vast majority (>90%) of associated genome loci for complex disease and traits fall in non-coding regions^4^. Hence, it is of exceptional utility to explore these datasets and derive novel hypothesis to interpret non-coding genome. Unlike the protein coding region where there is a clear genetic code, incorporating broader sequence context is critical to understand functional effects of regulatory variants, which requires more powerful and semantic-rich representational model to capture higher-order complexity in the region. Deep leaning has transformed ranges of tasks in computer vision and natural language processing (NLP). In bioinformatics field, deep learning based computational methods have also been proposed in various applications, such as predicting molecular phenotypes based on raw DNA sequence as input and achieved better performance than traditional machine learning approach, as referred in an excellent review paper^5^. One classical model is DeepSEA^6^, pioneering to apply deep convolutional neural network (CNN) architecture to extract features of genome sequence given 1,000-bp context and train on chromatin profiles in a supervised multitask joint learning manner. DeepSEA is able to predict the binary presence or absence of 919 chromatin marks.

The inherent sequential nature of genome is analogous to documents composed of words,characters and phrases. The exciting advance of NLP field has shed light on using similar strategy to extract general and transferable information from biological sequence. Neural network model was introduced into NLP since 2013. Word2vec^8^ was proposed to learn distributional vector embeddings of each word to capture their similarities given the sentence context. Word2vec uses multilayer perceptron (MLP)^7^ to predict neighboring words given center word (called skip-gram) or predict center word given neighboring words (called ‘Continuous Bag of Words’, CBOW). The learned word vectors can then be directly queried for downstream text classification tasks. Word2vec essentially relies on modelling co-occurrence probabilities without considering word position information and static embeddings cannot handle words with multiple meanings, so-called polysemous words. Traditional CNN-based feature extractors rely on local parameter sharing and the pooling operation may lead to loss of global information. Recurrent Neural Network (RNN)^9,10^ is an alternative architecture to process sequential data. RNN can capture position dependency information via passing the memory state from previous elements. RNN’s fundamental constraint of sequential operation leads to difficulty of parallelization and faces the risk of vanishing gradient when processing longer sequence. In 2017, Transformer^11^ has emerged as a powerful architecture that replies completely on attention mechanism to draw global dependencies in Seq2Seq modelling task. In the encoder part, self-attention mechanism relating different positions across a single sequence to compute a contextualized representation with better parallelization. Transformer can tackle long-range dependency without position bias, outperforming CNNs or RNNs in many global sequence classification tasks. On the other hand, CNN is better at capturing locality.

Since 2018, a new wave of pre-trained language models using self-supervision techniques has emerged as a core trend in NLP, including RNN-based ELMo^12^, ULMFiT^13^, Transformer-based OpenAI-GPT^14^ and Google-BERT^15^. Instead of conventional left-to-right unidirectional modeling, BERT, which stands for Bidirectional Encoder Representations from Transformers, leveraged a multilayer bidirectional Transformer architecture to pre-train on large unlabeled corpora by jointly incorporating both left and right context. The pre-trained model learns contextualized token embeddings through two proxy training objectives: MLM (Masked Language Model), predicting randomly masked tokens and NSP (Next Sentence Prediction), predicting whether two sentences follow each other. The pre-trained BERT can then be easily fine-tuned to various downstream NLP tasks and obtained new state-of-the-art results competing with human performance. Thereafter, a series of pre-trained model spring up to further improve performance, such as XLNet^16^, UniLM^17^, MASS^18^, MT-DNN^19^, XLM^20^, ALBERT^21^, RoBERTa^22^ and ELECTRA^23^. A comprehensive review can be found in an integrative reference^24^. One representative model, ALBERT, a lite version of BERT, establishes better result with significantly reduced model size through factorized embeddings and cross-layer parameter sharing techniques. Unlike model trained on general domain corpora, SciBERT^25^ and BioBERT^26^ are proposed based on BERT backbone and trained on large amount of multidisciplinary scientific literatures and biomedical text corpus, respectively. The domain-specific BioBERT achieves dramatic improvement in biomedical text-mining tasks. Recently, Transformer was reported by Facebook to learn protein structure and function^27^. DNABERT^28^ is recently proposed to learn human genome and composed of 12 Transformer layers with 768 hidden units and 12 attention heads in each layer, which is configured as a heavy version of BERT. However, features learned by Transformer are too general and not specific or sensitive to a single or few changes in the sequence, unable to satisfy the needs of interpreting human genome at base-resolution. Recently, Facebook and Google both propose introducing convolutional layer into Transformer architecture to bring soft inductive bias with better locality, namely ConViT^29^ and CoAtNets^30^.

Motivated by these observations, in this study we develop LOGO, a pre-trained language model with much lighter architecture than the pioneering DNABERT, which is composed of only 2 layers with 256 hidden units and 8 heads, to learn contextualized representations of reference genome hg19. LOGO with 3-mer tokenization contains around 1 million parameters while DNA-BERT contains 100 million parameters. LOGO shows substantially more effective parameter efficiency than DNABERT. To demonstrate the versatility of LOGO, we implement fine-tuning for multiple downstream tasks and obtain excellent performance from aspects of accuracy, speed, scalability, and robustness. Sequence-level classification tasks include promoter prediction, promoter-enhancer interaction prediction and chromatin features prediction. Another key innovation of LOGO is that we introduce a novel encoding scheme for alternative alleles and leverage a hybrid architecture by mixing convolution and self-attention to alleviate the locality-insensitivity issue of Transformer and facilitate functional prioritization of noncoding variants. DNABERT only reports high-attention variants by Transformer while LOGO leverages convolution and forces the model to see the nucleotide change with allelic-effects. We also propose a framework to embed prior knowledge into LOGO and explore better performance over original settings. LOGO provides a unified framework not only for sequence labelling or motif identification task, but also for SNP or indel prioritization with the goal of interpreting non-coding regions at base-resolution.

## Results

### LOGO learns contextualized representation of k-mers of human reference genome and achieves state-of-the-art performance in promoter prediction task

The backbone of LOGO processing flow is motivated by recently emerged Transformer-based bidirectional encoder model^15,21^ (Fig. 1a). We conduct pre-training on human reference genome hg19 comprised of totally 3 billion base pairs. We segment both forward and complementary chain of whole genome sequence into 100-bp bins and get 60 million segments. For each bin, we extend forward to 1,000-bp along the genome to create training instances, which are analogous to input sentences in the field of natural language.

**Fig.1.**
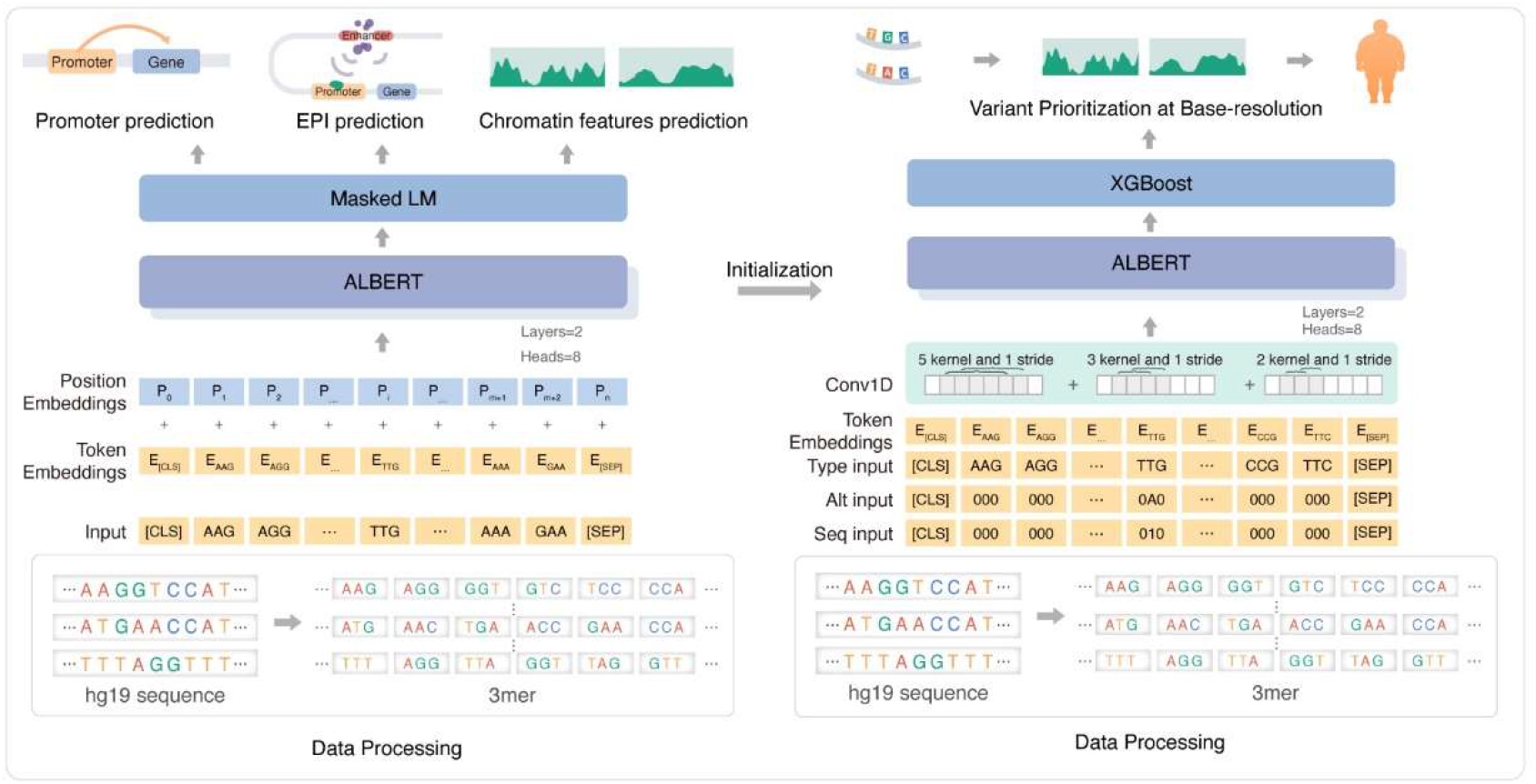
Overview of LOGO. LOGO is firstly pe-trained on human reference genome hg19 and then fine-tuned on several downstream tasks. LOGO uses self-attention based Transformer architecture, a light version language model (ALBERT) with only 2 self-attention layers with 256 hidden unites and 8 heads. Input genome sequence is represented by sequential k-mer vectors (k=3, 4,5,6) and then projected into a distributional embeddings. Token embeddings under 1,000-bpor 2,000-bpcontext are learned via masked language model (LM) task. The pre-training objective is to predict the randomly masked tokens by a Softmax layer over the vocabulary. Token embeddings and position embeddings are summed up and fed into ALBERT network. For sequence labelling task at global sequence level, including promoter identification, enhancer-promoter interaction (EPI) and chromatin features prediction, [CLS] token is used as global features extracted by LOGO, standing for aggregated representations of each input sequence for sequence classification task. [SEP] token stands for the end of each input sequence (Methods). For variant prioritization task, LOGO utilizes a multi-stream scheme to encode reference allele, alternative allele, and corresponding altered position as input. A convolutional layer is added to introduce locality to capture allelic-effect at base-resolution. Pre-trained LOGO weights are used as model initialization for variant prioritization task (Methods).

Conventional one-hot encoded representation for each nucleotide has limited vocabulary size of 5 characters (i.e., A, G, C, T, and Unknown/Undetected), which is considered as a semantically poor representation. K-mer encodes sequence into a certain length of successive nucleotides. For instance, a trinucleotide is a k-mer for which k=3. To increase the information content, we tokenize each sequence in the way of k-mer representation. The intuition is that each nucleotide is not independent such as codon rules in coding region and regulatory motifs in non-coding region. Recent phrase-level or entity-level masking strategy used by NLP community proved better performance. BERT or ALBERT generally allows maximum sequence length of 512 tokens, thus k-mer setting can dramatically reduce the number of tokens required to incorporating 1,000-bp context. Token vocabulary size equals to 5^k^ when using k-mer strategy. 7-mer or longer settings results in unbearable computation burden and memory overflow due to explosive vocabulary size. To balance the computation consumption and representation capacity, we evaluate four types of k-mer (k=3,4,5,6) to tokenize the genome. For any given sequence, different sets of k-mer representations can be generated when choosing different split positions. To avoid biased segmentation and further augment training data, we slide 1-bp (k-mer-1-stride) for each 1,000-bp sequence to generate multiple sets (n=k) of k-mer tokens as input training instances. (Tokenization details see Methods)

Before fed into the Transformer network, each token representation is created by summing its corresponding token and position embeddings. Token embeddings are learned through projecting the k-mer vectors into a distributional embedding space. To utilize the order of each sequence, we inject absolute position information of each input sequence and make the model learn position embeddings of the same size as token embeddings. During pre-training stage, we only adopt ‘masked language model’(MLM) task to train a bidirectional representation for human genome. 15% of tokens are randomly masked in each input sequence and the pre-training objective is to predict the masked token by a softmax layer over the vocabulary. We denote the k-mer tokens embedding size as E, the number of encoder layers as L, and the hidden layer embedding size as H, the number of self-attention heads as A. Visualization of model architecture can be seen in Fig. 1b. We pre-train LOGO with k-mer tokenization (k=3, 4, 5, 6) on hg19 for maximum 50 epochs on four Nvidia Tesla V100 32G GPU. Model hyperparameters are determined by choosing a model size as minimal as possible without compromising the performance. Detailed hyperparameters and pre-training configuration can be seen in Supplementary Table 1.

Bigger k leads to larger vocabulary size, therefore requiring increased model parameters, more memory usage and longer convergence time. We assess the pre-training performance based on the accuracy (ACC) of masked tokens prediction. 3-mer tokenization achieves the highest MLM accuracy with a minimum training time spent per epoch (Fig. 2a,2b). For all k-mer settings, LOGO can achieve inflection point of pre-training accuracy after 5 epochs, though already surpass 0.875 when training less than one epoch, revealing recurring sequence patterns of human genome is effectively learned. One epoch training time for 3-mer tokenization setting is around 11.4 hours, and we reach accuracy plateau (ACC=0.893) after 15 epochs. One epoch training time for 6-mer tokenization setting is around 70.8 hours, and we reach accuracy plateau (ACC=0.887) after 25 epochs. However, accuracy at pre-training stage is not directly correlated with utility of specific fine-tuning task. We assess all four k-mer settings for different downstream tasks and only report the best one. Other details of pre-training assessment can be found in Supplementary Table 2.

**Fig2.**
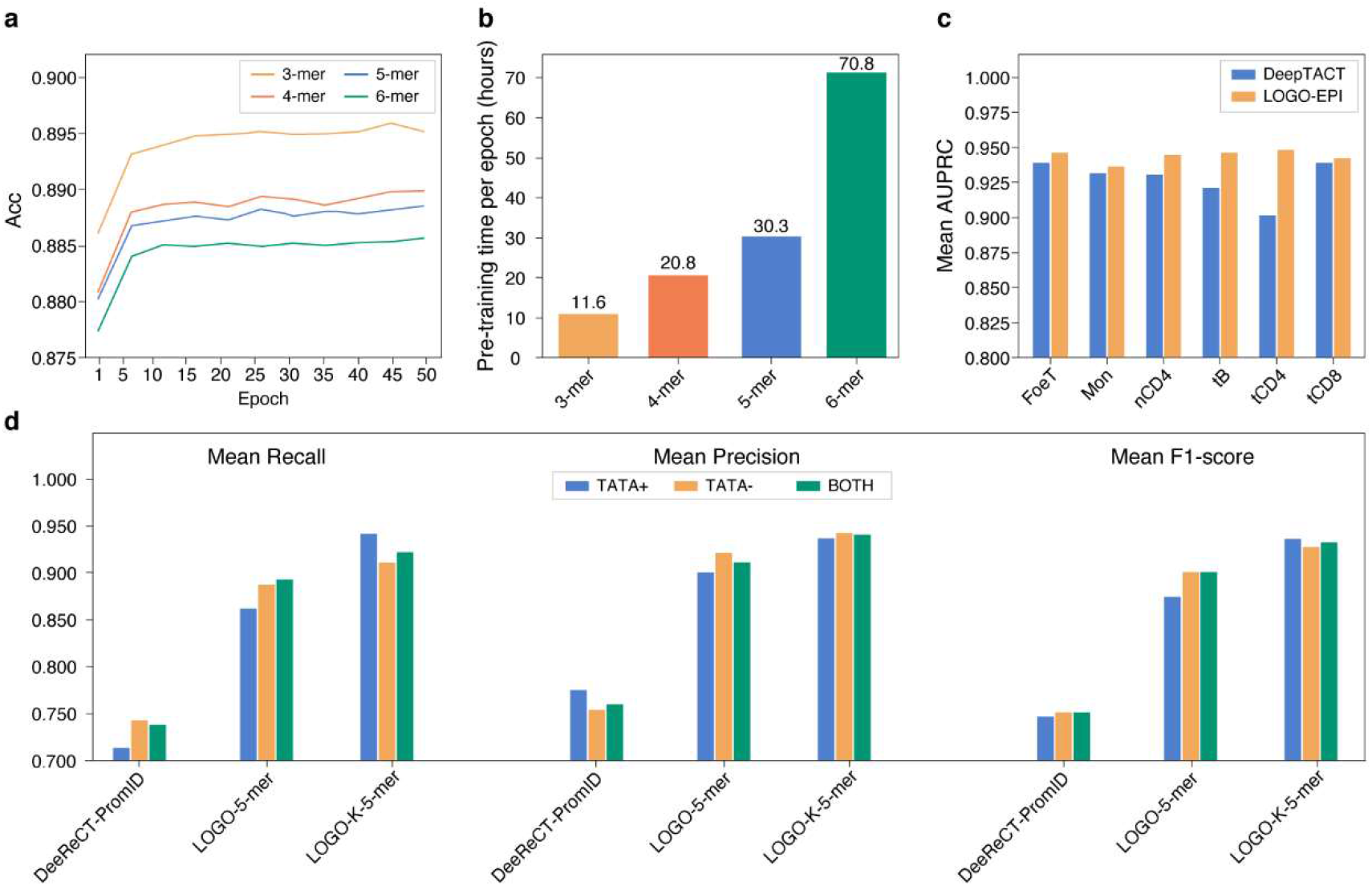
LOGO learns contextualized representation of k-mers of human reference genome and achieves state-of-the-art performance for promoter prediction and enhancer-promoter interaction prediction. **a**, Pre-training accuracy (ACC) plateaus after 5 epochs for all k-mer settings. 3-mer tokenization achieves the highest ACC. **b**, LOGO pre-training time of one epoch for all k-mer settings is plotted. (All settings are trained on four Nvidia Tesla V100 32G GPU). Larger k leads to longer training time due to larger vocabulary size. **c**, Pre-trained LOGO using 5-mer tokenization (LOGO-5-mer) is fine-tuned on promoter prediction task and evaluated against DeeReCT-PromID on promoter sequences from EPDnew Database, including ones with TATA-box(TATA+, n= 2,067), without TATA-box (TATA-, n=14,388) and jointly (Both, n=16,455). Knowledge embedded LOGO (LOGO-K-5-mer) further boost the performance. 11 annotations terms from GenBank, i.e., ‘CDS’, ‘exon’, ‘enhancer’, ‘insulator’, ‘conserved_region’, ‘protein_binding_site’, ‘pseudogene’, ‘DNAsel_hypersensitive_site’, ‘nucleotide_cleavage_site’, ‘silencer’ and ‘gene’ are introduced in one-hot encoded format as knowledge input. Metrics of mean Recall, mean Precision and mean F1-score are evaluated using 10-fold cross-validation. **d**, Pre-trained LOGO is fine-tuned on enhancer-promoter interaction prediction task (LOGO-EPI) and evaluated against DeepTACT on promoter capture Hi-C (PCHi-C) datasets in six different cell types, including fetal thymus (FoeT, n= 6,676), monocytes (Mon, n=8,062), naïve CD4+ T cell (nCD4, n=8,712), total B cells (tB, n=9,036), total CD4+ T cell (tCD4, n=8,282) and total COB+ T cell (tCD8, n=8,140). Mean area under precision-recall curve (AUPRC) are evaluated using 10-fold cross-validation.

We first evaluate the utility of pre-trained LOGO on human promoter prediction task via fine-tuning. Computational identification of promoters is analogous to sequence labeling or sentence classification task in NLP. Umarov et al. has developed CNN-based deep leaning models, DeeReCT-PromID^31^, to predict human RNA pol II promoters, outperforming other previous prediction tools. For benchmark purpose, we generate datasets in the same way with DeeReCT-PromID and define positive promoter region from -200 bp to +400 bp window around all potential Transcription Starting Site (TSS) from EPDnew Database^32^. Promoters with TATA-box (TATA+) and without TATA-box (TATA-) are assessed separately and afterwards jointly (Both), which leads to 2,067, 14,388 and 16,455 positive sequences, respectively. Negative ones are constructed by randomly sampling outside the promoter region without containing a known TSS. Leveraging previously pre-trained LOGO model weights as initialization, we simply plug in these 600-bp sequence inputs, tokenize them via different k-mer-1-stride settings (k=3, 4, 5, 6) and feed them into LOGO. When conducting sequence classification task, model input starts with a token [CLS] as in BERT and ALBERT. We use the final hidden vector of [CLS] token as the aggregated representation for classification tasks and fine-tune model parameters in an end-to-end manner, which only introducing few extra parameters in the final classification layer with sigmoid outputs. We use batch size of 128, set early-stop rules and fine-tune the model at most 20 epochs. The average training time per epoch is only around 45 seconds, which demonstrates excellent efficiency of ‘pre-training and fine-tuning’ paradigm. The best hyper-parameters are chosen based on the validation sets (Methods). We evaluate different k-mer settings of LOGO on promoter prediction tasks. LOGO significantly outperforms DeeReCT-PromID in all-settings as evaluated by Precision, Recall and F1-score metrics, as shown in Fig 2d and Supplementary Table 3. LOGO with 5-mer setting (LOGO-5-mer) achieves 15.0% point absolute improvement of mean F1-score than CNN-based DeeReCT-PromID (10-fold cross-validation) in case ‘Both’. LOGO has demonstrated its powerful representation utility, which suggests bidirectional attention-based architecture confers an advantage to capture complex semantics of promoter structure than CNN-based model.

We further explore a framework to integrate prior knowledge into LOGO on promoter prediction task. GenBank^33^ contains rich functional annotations of human genome sequence, including CDS, exon, gene, promoter, enhancer, silencer, pseudogene, insulator, conserved region, protein binding site, DNAse I hypersensitive site, nucleotide cleavage site and so on. These annotations can be regarded as prior knowledge of sequence inputs. We download 11 annotations terms from GenBank, i.e., ‘CDS’, ‘exon’, ‘enhancer’, ‘insulator’, ‘conserved_region’, ‘protein_binding_site’, ‘pseudogene’, ‘DNAseI_hypersensitive_site’, ‘nucleotide_cleavage_site’, ‘silencer’ and ‘gene’. Annotations of ‘promoter’ are abandoned to avoid direct label leakage. We generate annotation labels for each input sequence in a start-to-end spanning mode based on hg19 coordinate. We propose a knowledge-enabled LOGO by adding input layers of one-hot encoded annotations and concatenate with k-mer input (Supplementary Figure 1). Knowledge embeddings, position embeddings and token embeddings are summed up and then fed into Transformer network for fine-tuning tasks in an end-to-end manner (Fig. 1b). Knowledge embedded LOGO with 5-mer setting (LOGO-K-5-mer) achieves F1-score of 0.933, yielding extra 3.2% absolute improvement than LOGO-5-mer (Fig. 2d). We demonstrate the configurability and utility of knowledge-embedded framework for genome sequence labelling. We caution that this attempt is preliminary and might introduce position bias or indirect label leakage into the model. We envision that rationally incorporating experimentally validated human knowledge can assist in developing better sequence representation model for scientific discovery.

### LOGO can be used to predict regulatory interactions between enhancer-promoter sequence pairs

Predicting 3D chromatin contacts between promoters and enhancers is critical to understand transcriptional regulation in specific cell-lines or tissues. Computational approach is needed to improve the resolution of Hi-C data and detect genome-wide physical interactions at corresponding regulatory elements. This task is analogous to general inter-sentence modelling in NLP, such as sentence pairs in paraphrasing, hypothesis-premise pairs in entailment, and question-passage pairs in question-answering task. We draw lessons from NLP field and consider 3D chromatin contacts prediction as a sequence pairing problem.

Li et al. proposed DeepTACT^34^, a CNN and RNN mixed deep learning model with one attention layer to predict enhancer-promoter interactions (EPI). DeepTACT leverage both raw sequence and chromatin accessibility information, but we only benchmark LOGO against DeepTACT version without accessibility information input due to unavailability of processed chromatin features. We retrain DeepTACT and fine-tune LOGO (LOGO-EPI) on the same bootstrapped dataset provided by the author of DeepTACT, which contains three parts: 2000-bp window enhancer sequences^35^, 1000-bp window promoter sequences^36^ and paired enhancer-promoter interaction (EPI) labels from promoter capture Hi-C (PCHi-C) experiments in six different cell types^37^, i.e., fetal thymus (FoeT), monocytes (Mon), naïve CD4+ T cell (nCD4), total B cells (tB), total CD4+ T cell (tCD4) and total CD8+ T cell (tCD8) (Supplementary Table 4). Similar data augmentation technique is applied to generate larger positive training examples. The performance of each model is evaluated by tenfold cross-validation. LOGO-EPI uses 6-mer setting to tokenize input sequences. We add one 1D convolution operation for each input promoter or enhancer sequence before feeding into Transformer network. The underlying intuition is to avoid large fluctuations of token embeddings during fine-tuning stage and ensure certain disparities among tokens (Supplementary Figure 2). The learned representations of paired promoter and enhancer sequence are concatenated and fed into the final binary classification layer. (Model architecture seen in Fig. S1). LOGO-EPI achieve 0.23-4.47% absolute improvement than DeepTACT on AUPRC for six cell lines (Fig. 2c). LOGO-EPI outperforms DeepTACT most significantly for tCD4 and LOGO-EPI yields more consistent performance while DeepTACT fluctuates across different cell lines (AUPRC details in Supplementary Table 5).

### LOGO achieves superior performance on chromatin features prediction with significantly reduced model size and much less training time than previous models

Next, we move on to compare LOGO against CNN-based DeepSEA^6^ to predict chromatin features from DNA sequence. Unlike fully supervised training manner as DeepSEA, we fine-tune the pre-trained LOGO with chromatin features prediction task and demonstrate higher accuracy with significantly improved scalability. To make proper comparison as well as demonstrate model scalability, we use three sets of chromatin profiles with some overlaps; the first one is the same as original DeepSEA paper with 919 chromatin features, the second one is 2,002 chromatin features expanded by DeepSEA developer group reported in ExPecto^38^, and we construct the third one of 3,357 chromatin features by integrating ExPecto’s 690 transcriptional factors (TF) binding features with recently released 2,850 EpiMap^39^ (for epigenome integration across multiple annotation projects) features after deduplication. (Data details in Methods)

In the first task, we use the same training, validation, and test sets as in DeepSEA. LOGO-919 obtains 0.70% 0.70% and 0.80% absolute improvement of median AUROC than downloaded DeepSEA for predicting 690 TF binding, 125 DNase hypersensitive sites (DHSs) and 104 histone modification marks (HM), respectively (Fig. 3a). The maximum increase is for transcription repressor ZNF274 binding in HepG2 cell line (AUROC=0.703 by LOGO-919 versus AUROC =0.582 by DeepSEA). LOGO’s model architecture and training strategy confer huge advantage on computation efficiency and memory consumption over traditional deep CNN-based architecture completely trained in a multitask supervised manner. LOGO has much smaller parameter size compared to DeepSEA. LOGO-919 contains around 1.52 million parameters, which is 34x fewer than DeepSEA’s 52.8 million parameters (Fig. 3c). LOGO-919 obtains better performance than downloaded DeepSEA after 33 hours of pre-training and fine-tuning on 4 Nvidia Tesla V100 GPU (around 110 hours on 1 Nvidia TITAN Xp Pascal GPU). We also retrain DeepSEA from scratch on 1 Nvidia Titan Pascal GPU and stop after 1 month (720 hours) and reproduce slightly poorer performance than the downloaded version, which indicates LOGO-919 takes at least 6x shorter training time than DeepSEA. (Fig. 3d). The improvement in parameter efficiency is the most critical advantage of LOGO framework, which gives LOGO superior advantage to extend to ever-growing chromatin maps. The learned semantic-rich representation for k-mer tokens in a self-supervised manner alleviate the excessive needs of cumbersome fitting for multiple supervised tasks from scratch. To demonstrate this concept, we conduct the second experiment using 2,002 chromatin features, including 690 TF binding, 334 DHSs and 978 HM features as reported in ExPecto model (Details in Supplementary Table 9). ExPecto also used CNN-based architecture and extend DeepSEA by doubling the number of convolution layers to increase model depth to satisfy doubled learning objectives, ending up with around 150 million parameters, nearly 3-fold of DeepSEA. We retrain the chromatin marks prediction part of ExPecto and stop after 1000 hours. The learning tasks double and the model parameters triple than DeepSEA, which shows severe lack of scalability. For fair comparison against ExPecto, we incorporate 2,000-bp context window while remaining other settings unchanged and fine-tune LOGO-2002 within 66 hours. We choose the time upper bound according to the intuition of doubled training time (66 hours vs 33 hours) for doubled tasks (2,002 features vs 919 features) (Fig. 3d). The model size only marginally increases (1.87 million parameters) due to longer input context and additional parameters of final classification layer (Fig. 3c). The model backbone remains unchanged, and the results show LOGO-2002 can achieve comparable median AUROC with retrained ExPecto on held-out chromosome within 66 hours. The median AUROC for TF, DHSs and HM is 0.954, 0.913, 0.883 respectively (Fig. 3b). We demonstrate that LOGO can scale easily via pre-training and fine-tuning paradigm with benefits of computational speed and reduced parameterization. It is noted that DNABERT contains more than 100 million parameters while LOGO only contains 1 million parameter, which demonstrating LOGO’s superior efficiency of parameter sharing among attention layers. In addition, LOGO predicts chromatin features in a jointly multi-task manner while DNABERT only supports TF-binding site prediction one TF by one TF, which is considered as a much simpler task and cannot effectively transfer the knowledge among different chromatin annotations.

**Fig3.**
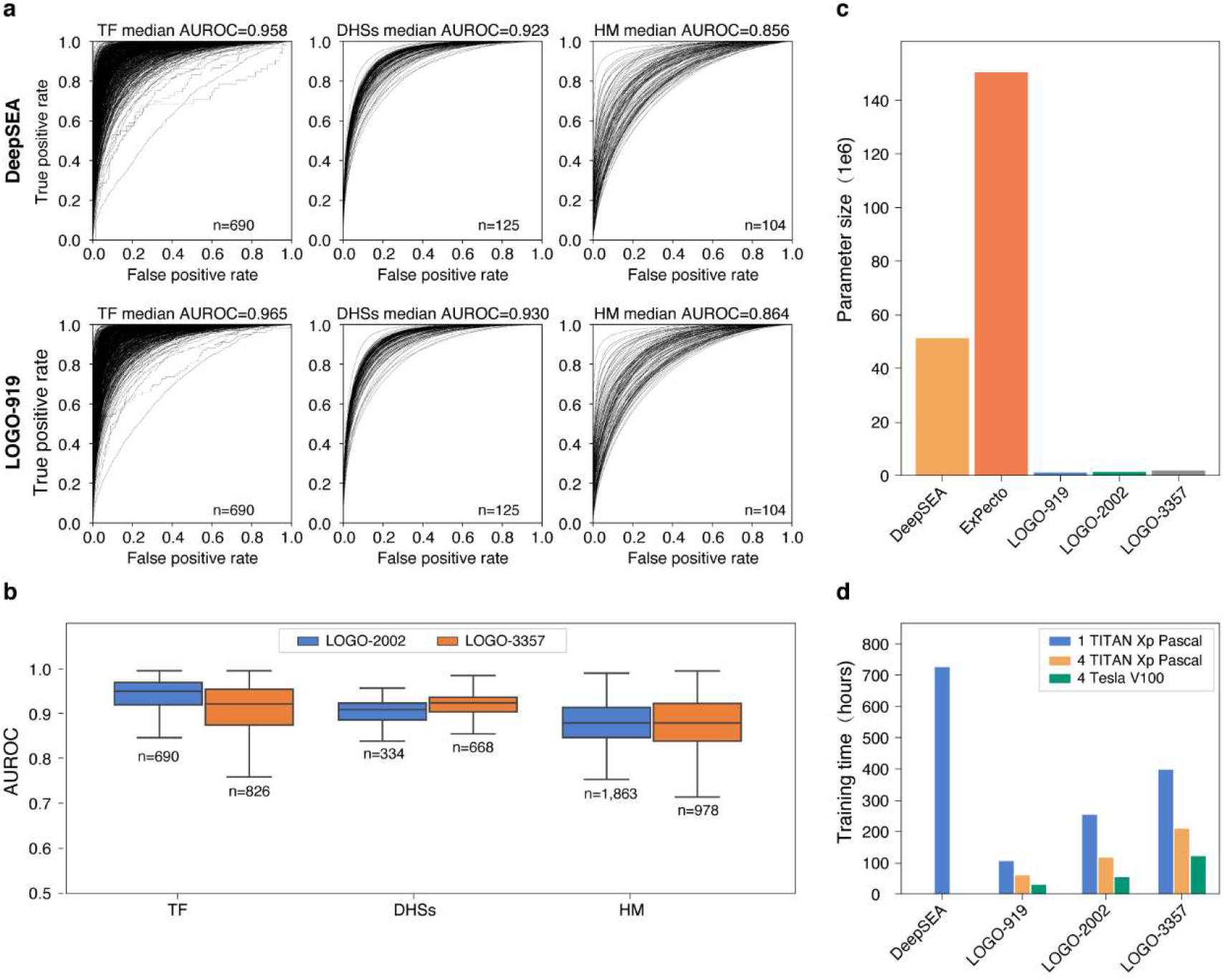
LOGO line-tuned on chromatin profiles outperforms DeepSEA with significantly reduced parameter size and consumes much less training time. **a**, Receiver operating characteristic (ROC) curve is plotted to compare predictive power between DeepSEA (top) and LOGO-919 (down) for 690 Transcription factor binding (TF), 125 DNase hypersensitive sites (DHSs) and 104 histone modification marks (HM) on held-out chromosome using 1,000-bp context window. Metrics of median area under receiver operating curve (AUROC) for all TF, DHSs and HM are displayed above each curve. **b**, Boxplot shows AUROC for three types of features predicted by LOGO-2002 (Pretrained LOGO with 2,000-bp context fine-tuned on 2,002 chromatin fea tures from ExPecto, n= 690 TF, 334 DHSs and 978 HM, respectively) and LOGO-3357 (Pretrained LOGO with 2,000-bp context fine-tuned on 3,357 chromatin leatures after integrating EpiMap features with ExPecto features, n= 826 TF, 668 DHSs and 1,863 HM, respectively) (Methods). Box plots show median, upper, and lower quartiles, and highest and lowest values excluding outliers. **c**, Plot shows parameter size of DeepSEA, ExPecto, LOGO-919, LOGO-2002 and LOGO-3357. **d**, Plot shows compari son of training time among DeepSEA, LOGO-919, LOGO-2002 and LOGO-3357. Training time for DeepSEA is recorded as duration of reproducing DeepSEA from scratch using 1 TITAN Xp Pascal GPU, with slightly lower model performance than downloaded version. Training time of LOGO-919, LOGO-2002 and LOGO-3357 include both pre-training (1-2 epochs) and fine-tuning. Three sets of GPU configuration used to fine-tune LOGO are indicated by different colors.

Incorporating more comprehensive chromatin profiles or using task-specific features have both been reported useful for functional analysis of noncoding variants^62,63^. Abundant experimental mappings of human epigenomes are continuously accumulating chromatin profiles for more cell types and tissues. Further expanded chromatin features require larger CNN-based models with explosive parameters, while LOGO can be easily extended to more chromatin features with marginally increased parameters. LOGO demonstrates its powerful scalability and easy deployment, which is of critical importance to tackle even larger-scale functional map. To further prove this concept, in the third experiment, we utilize the most comprehensive chromatin profiles from EpiMap and integrate with all TF features from ExPecto. We construct datasets of 2,000-bp context window paired with a label vector for 3357 chromatin features using Selene^40^. We fine-tune LOGO-3357 using the same model architecture as LOGO-2002, achieving median AUROC of 0.926, 0.928, 0.883 for 826 TF, 668 DHSs and 1,863 HM features respectively (Fig. 3b). The slightly lower performance than LOGO-919 and LOGO-2002 is mainly due to less stringent dataset construction and less precise position calibration for EpiMap related features. LOGO-3357 has nearly 2.22 million parameters, which is still significantly fewer than DeepSEA and ExPecto (Fig. 3c), again demonstrating its scalability to triple chromatin features without the need of increasing model size substantially or extending disproportionate training time. All training details can be found in Supplementary Table 6.

### LOGO can be used to predict functional effects of noncoding variant at base-resolution and provides mechanistic insights for investigating complex disease

The associated loci identified by GWAS provide abundant information regarding the genetic basis of human complex diseases and traits. Nevertheless, owing to Linkage Disequilibrium (LD), it remains challenging to identify high-resolution causal variants in an interpretable manner^41^. Variants from GWAS catalog predominantly consists of marginally associated variants that have not been fine-mapped. We attempt to extend LOGO to prioritize noncoding functional variants for complex disease based on the predicted signals of the above three sets of chromatin features. We anticipate that, if a complex disease-related variant exerts its effect via disruption of TF binding motif or via alteration of DNA accessibility or histone modification, this SNP can be identified *de novo* from sequence by LOGO. We choose type-2 diabetes (T2D) as an example to test this hypothesis and construct evaluation datasets from the latest published literatures and GWAS resources. (Data Details see in Methods). We employ the same DeepSEA E-value metric to estimate the regulatory potential of a SNP by comparing the allele-specific probabilities per SNP to one million random SNPs from the 1,000 Genome Project (Phase 3).

First, we demonstrate LOGO can be used to prioritize putative causal regulatory variants from GWAS reported T2D-associated loci. We hypothesize that if LOGO is fine-tuned on more comprehensive chromatin profiles, it can identify more functional variants within LD blocks. We download all T2D-associated SNPs from GWAS Catalog^42^ (2020-05-14 version, p-value ranging from 9×10^−6^ to 6×10^−447^) and corresponding LD SNPs (r^2^>0.2) from LDlink^43^, resulting in 156,175 SNPs after deduplication. A variant is considered as functional significant if at least one chromatin feature’s E-value is equal or less than 1×10^−5 6,44^. LOGO-3357 can identify more functional SNPs (n=14,764) than LOGO-2002 (n=729) and LOGO-919 (n=374) (Supplementary Table 7). Within the 71 GWAS Catalog lead SNPs identified by LOGO-3357, 30 of them reach genome-wide significant (p-value<5×10^−8^) in at least one GWAS. We divide all functional significant SNPs identified by LOGO-2002 and LOGO-3357 into two groups (r^2^ ≥0.5 and r^2^<0.5), we compare mean activated chromatin features (E-value<1×10^−5^) of each group and discover that SNPs with higher LD activate more chromatin marks (p-value=0.00093 for LOGO 3357, p-value=0.02 for LOGO 2002, by Mann Whitney U-Test, Supplementary Figure 3). We demonstrate that LOGO has the potential of fine-mapping causal variants within LD block in an explainable manner and the model fine-tuned on more chromatin features provides more functional attributions. We further conduct tissue-enrichment analysis for all putative functional variants identified by LOGO-2002. Hypergeometric test is used to evaluate whether activated chromatin features are enriched in certain categories. We find that these SNPs are functionally enriched in 18 categories out of total 27 with activation signals, including smooth muscle(n=51), lymphoblastoid(n=45), adipose(n=20), muscle(n=63), spleen(n=10), and liver(n=48), (-log(p-value) = 11.1, 5.3, 5.0, 3.2, 3.1, and 1.3 respectively), which is consistent with years of pathogenesis research that insulin mainly acts on liver, muscle and adipose as T2D-relevant tissues (Fig. 4a, Supplementary Figure 8). Recent integrative epigenomics study^39^ (EpiMap) leverages enhancer sharing tree to investigate tissue enrichment of T2D-related SNPs and indicates that T2D is polyfactorial traits enriched in up to 18 tissue categories out of 33 tested. Sequence-based LOGO-2002 shows similar diversity of enrichment with significant tissue overlaps. Islets only constitute ∼1% of the pancreas and specific annotations of islet epigenome are absent in ENCODE and Roadmap Epigenomes Project^45^. Thus, analyzing pancreas organ alone fails to provide reliable information of islet epigenomes. Thurner^45^ and Varshney^46^ have specifically annotated promoter/enhancer state of islets, which are regarded as typical T2D relevant cell types. Specifically, 10 functional SNPs identified by LOGO-3357 are overlapping with islets-specific promoter/enhancer state with at least 5 activated chromatin features (E-value<1×10^−5^) (Table 1). The disruption of regulatory function is consistent with previous report that parts of T2D-related risk variants are considered to act through primary effects on beta-cell function. For instance, T2D-risk allele at rs9693089 (FAM167A locus) locates at the active enhancer state of islet sample identified by Varshney^46^ and has been reported to be associated with very low-density lipoprotein (VLDL) synthesis by Kraja^47^. The corresponding activated chromatin features include H3K4me3, H3K4me1, H3K27ac. Even though LOGO-3357 is not specifically trained on islet chromatin marks, this experiment demonstrates that deep learning based methods have the potential of providing extra informativeness using sequence alone as input^48^.

**Fig4.**
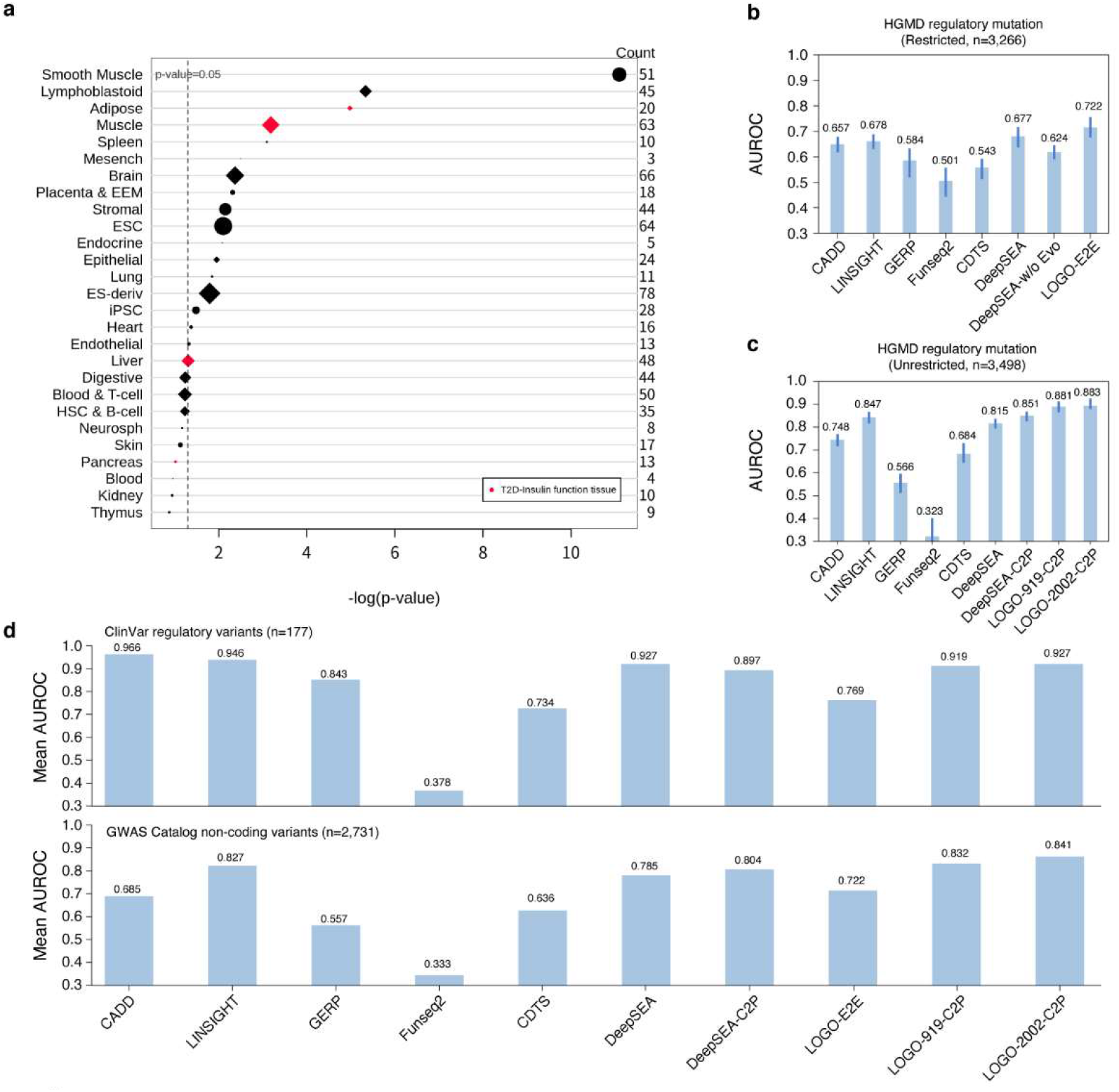
LOGO can be used to infer underlying regulatory mechanisms of T2D GWAS signals and prioritize functional variants for both inherited disease and complex trait or disease. **a**, Tissue enrichment results for significant T2D-related variants identified by LOGO-2002 located in promoter/enhancer state regions, sorted by -log(p-value)(Hypergeometric test). The size of circle/diamond represents the number of activated chromatin marks in corresponding tissue or cell types, also displayedon the right side of the plot. Red symbols indicate 4 well-known tissue types related with T2D. Tissue or cell types enriched by EpiMap for T2D GWAS signals are represented by diamond shape. T2D, type 2 diabetes; EEM, extra-embryonic membranes; ESC, embryonic stem cell; ES-deriv, embryonic stem-derived; iPSC, induced pluripotent stem cell; HSC, hematopoietic stem cell. **b**, Prediction power of various models for prioritizing HGMD regulatory mutations (n=3,266) against negative controls (n=3,266) in restricted scenario that negative samples matched to positive ones within 1 kb. **c**, Prediction power of various models for prioritizing HGMD regulatory variants (n=3,498) against negative controls (n=3,690) in unrestricted scenario of random sampling. Mean AUROC of 10-fold cross validation for b and c are reported, Error bars represent standard deviations. **d**, Comparison of model performance by metric of mean AUROC on held-out test set from inherited disease domain (top: n=177 stringent ClinVar regulatory variants, negative controls are bootstrapped 10 times) and complex trait or disease domain (bottom: n=2,731 genome-widesignificant non-coding variants from GWAS Catalog, positive variants are bootstrapped 10 times). Analysis with LOGO-E2E, LOGO-919-C2P and LOGO-2002-C2P are compared with that of CADD, LINSIGHT, GERP, Funseq2, COTS, DeepSEA and DeepSEA-C2P. DeepSEA and DeepSEA-w/o Evo means DeepSEA derived functional significant score including and excluding 4 evolutionary features, respectively. DeepSEA-C2P represents a logistic regression model based on 919 chromatin marks and 4 evolutionary features. LOGO-E2E means LOGO fine-tuned on binary label of allelic deleteriousness in an end-to-end manner. LOGO-919-C2P and LOGO-2002-C2P stand for LOGO fine-tuned on prediction of 919/2002 chromatin features with 4 evolutionary features followed by allele-specific variant prioritization via logistic regression model.

**Table 1.** Significant SNPs identified by LOGO-3357 overlapped with islet promoter/enhancer regions^a^

| RS ID | Locus | Significant Marks<sup>b</sup> | min E-value<sup>c</sup> | GWAS P-value<sup>d</sup> | GWAS Odds<sup>d</sup> | 1000G AF | Paper (PMID) |
| --- | --- | --- | --- | --- | --- | --- | --- |
| rs9693089 | *FAM167A* | H3K4me3 | 0.000001 | - | - | 0.68111 | 23192668 |
| chr8:11298385 A-G |  | H3K4me1 |  |  |  |  |  |
|  |  | H3K27ac |  |  |  |  |  |
| rs4735337 | *TP53INP1* | H3K4me3 | 0.000001 | - | - | 0.552516 | 25393876 |
| chr8:95973465 T-C |  | H3K4me1 |  |  |  |  |  |
|  |  | DNase-seq |  |  |  |  |  |
| rs1126899 | *CPA1* | H3K4me1 | 0.000001 | - | - | 0.582268 | - |
| chr7:130021488 G-C |  |  |  |  |  |  |  |
| rs11774700 | *LOC105375716* | HNF4G | 0.000001 | - | - | 0.271965 | 21188353 |
| chr8:118220270 T-C |  |  |  |  |  |  |  |
| rs163800 | *CTSZ* | H3K27ac | 0.000001 | - | - | 0.000399361 | 29795304 |
| chr20:57578508 T-C |  |  |  |  |  |  |  |
| rs4383259 | *ZNF347* | H3K4me1 | 0.000001 | - | - | 0.752196 | 24306210 |
| chr19:53661337 A-G |  | H3K27ac |  |  |  |  |  |
| rs3176447 | *CDKN2C* | DNase-seq | 0.000001 | - | - | 0.0740815 | 21145615 |
| chr1:51433687 T-A |  |  |  |  |  |  |  |
| rs11671664 | *GIPR* | H3K27me3 | 0.000001 | 3E-12 | 4.22[2.73-5.71] | 0.155152 | 27480816 |
| chr19:46172278 G-A |  |  |  |  |  |  |  |
| rs998451 | *TMEM163* | CEBPB | 0.000001 | - | - | 0.10643 | 24843659 |
| chr2:135429288 G-A |  |  |  |  |  |  |  |
| rs1776897 | - | EP300 | 0.000004 | - | - | 0.776757 | 27104953 |
| chr6:34195011 G-T |  |  |  |  |  |  |  |

<sup>a</sup>Islet promoter and enhancer regions are annotated by Thurner<sup>45</sup> and Varshney<sup>46</sup>. <sup>b</sup>Significant Marks means all chromatin marks with E-value<1×10<sup>-5</sup>. <sup>c</sup>Min E-value means the minimum E-value of corresponding chromatin mark activated by LOGO-3357. <sup>d</sup>GWAS P-value and GWAS odds are only shown for lead SNP reported from GWAS Catalog.

Second, we demonstrate that LOGO can be a sequence-based tool to help interpret those GWAS signals with possible population bias or sample size limitations. The statistic power of GWAS relies heavily on sample size, allele frequency and effect size of candidate SNPs^49^. GWAS with inadequate sample size can result in a multitude of nominally significant loci (p-value < 0.05). This problem is mainly mitigated by expanding sample size or conducting meta-analysis across cohorts or ethnic groups. Sequence based LOGO model is expected not to be affected by allele frequency or population bias and can evaluate both common and rare variants *ab initio*. We illustrate this potential using following examples. In study GCST005414^50^, rs340874 (PROX1 locus) reaches nominally significant (p-value=1 × 10^−7^). However, this SNP achieves genome-wide significant in study GCST009379^51^/GCST006867^52^ (p-value=2×10^−22^ and 8×10^−18^, respectively) with larger sample size. PROX1 has been reported to be associated with after-meal metabolism^53^, non-esterified fatty acids, and glucose metabolism^54^, which is also validated in both Japanese and Chinese populations^55,56^ (MAF=0.376). LOGO-3357 can directly identify rs340874 as a functional significant SNP (1 activated feature with E-value≤1×10^−5^, transcription factor POLR2A). PROX1 is reported to be a target gene of the POLR2A transcription factor from the ENCODE Transcription Factor Targets dataset. Rs896854 (TP53INP1 locus) is perceived to be associated with lipid levels of Chinese population with nominal significant signal (p-value=2×10^−6^) in study GCST004894^57^ and genome-wide significant signal (p-value=1×10^−9^) in study GCST000712^58^. Rs516946 (ANK1 locus) is reported in several independent studies to be correlated with decrease of insulin level and dysfunction of pancreatic islet cells at nearby site^59^. Again, LOGO-3357 can identify both rs896854 (1 activated feature with E-value≤1×10^−5^, Blood & T-cell, H3K9me3) and rs516946 (1 activated feature with E-value≤1×10^−5^, Other, DNase-seq) to be functional. Furthermore, we evaluate another 43 regulatory variants with posterior probability of association (PPA) >80% in a recent fine-mapping study^60^. 2 SNPs are identified functional significant by LOGO-3357 (rs340874 at PROX1 locus and rs76549217 at ANKH locus). Another largest-scale T2D meta-analysis study accumulates 1.4 million samples and discovered 318 new loci^61^, out of which LOGO-3357 can identify 14 SNPs to be functional. It is worth mentioning that all these 14 SNPs do not reach genome-wide significance in other populations except European ancestry. This result further indicates the unbiased predictive power of LOGO. 16 reported SNPs validated by LOGO-3357 with corresponding activated features are listed in Supplementary Table 8. We demonstrate that LOGO fine-tuned on chromatin features can help interpret GWAS non-coding SNPs and provide hints regarding underlying tissue-specific regulatory mechanism.

### Introducing locality-sensitive encoding scheme and convolution facilitates prioritizing functional variants for both inherited disease and complex trait or disease

Next, we move on evaluate whether fine-tuning LOGO can be used to develop functional predictor of pathogenic regulatory single-nucleotide variants (SNV) or common GWAS phenotype-associated SNPs. We define two schemes of fine-tuning: end-to-end training on binary label of deleteriousness (LOGO-E2E) and two-stage training of chromatin features prediction followed by variant prioritization (LOGO-C2P)^6,62^. Perturbation of molecular phenotypes can serve as an indicator of potential deleteriousness inspired by DeepSEA. We compare LOGO against 6 common predictors, including evolution-based method (GERP)^64^, sequence-based predictor based on chromatin effect signals with 4 evolutionary conservation features (DeepSEA)^6^, functional genome-based method (Funseq2)^65^, evolutionary method incorporating functional genome features (LINSIGHT)^66^, machine learning based classifier (CADD)^67^ and genome diversity metric (CDTS)^68^. It is noted that DeepSEA and CADD can provide allele-specific evaluations, whereas others assign identical scores to all alternative variants. Our predictors, LOGO-E2E and LOGO-C2P, are designed to capture allelic effect.

For variants associated with inherited human diseases, we extract a dataset from Human Gene Mutation Database (version 2019-03)^69^ to define positive examples of stringent regulatory mutations. We construct negative controls from 1000 Genomes Project^70^ SNPs by stringent frequency and population control, resulting in 3,498 pathogenic regulatory mutations and 3,690 negatives (total 7,188 variants, Supplementary Figure 6). We use 10-fold cross-validation to make robust comparison. For each fold, test variants are ensured to be scorable across methods. To increase stringency, we consider two schemes of negative sets selection: random sampling (unrestricted), and negative samples matched to positive ones within 1 kb (restricted, total 6,532 variants). Dataset construction details are illustrated in Methods.

For LOGO-E2E, we use three layers to encode variant presence and allelic information at specific position, including Ref layer, Alt layer and Variant Type layer. Ref layer is used to encode 1,000-bp context with 6-mer-1-stride setting. (Model architecture in Supplementary Figure 4) Alt layer is used to encode alternative allele at certain position to enforce the model to see directional alteration. Another Variant Type layer is set as default for SNV. By this means, we explicitly encode the alternative allele, ensure the ALT allele is always the effect allele. Each variant with surrounding context of certain length will be encoded as a matrix input containing ‘Ref’, ‘Alt’ and ‘Type’ information. 1-dimension convolutional layer is added before feeding token embeddings into LOGO to learn the binary deleteriousness effect of target variant. Fine-tuning LOGO in this way is expected to learn allelic pathogenicity. For LOGO-C2P, we follow similar pipelines in DeepSEA’s functional SNP prioritization part and firstly use previously trained LOGO-919/LOGO-2002 to generate chromatin effect features for both reference and alternative allele. We then conduct the same absolute difference and relative log fold change transformation as DeepSEA and feed these features into boosted logistic regression model to train the classifier at the second stage. It is worth mentioning that we discard z-score transformation used in DeepSEA-C2P classifier. We also assess the difference between preserving or removing evolutionary conservation features. The original scores of LINSIGHT, CADD, FunSeq2, GERP, CDTS and DeepSEA functional significant score are used to obtain the binary classification result with full range of thresholds.

In end-to-end setting, LOGO-E2E outperforms all other methods in restricted negative control scenario (AUROC=0.722) (Fig. 4b) and performs the second in the scenario of unrestricted negative control (AUROC=0.823). It is consistent with previous finding that restricted scenario poses more difficulties for distinguishing functional sites from surroundings than separating functional regions from genome background. Nonetheless, LOGO-E2E leverages an Alt token layer to enforce the model to explicitly encode allele position and directional mutation event to be distinguished from nearby unchanged context, which equips the model with allelic specificity under 1,000-bp context. For the less challenging unrestricted task, LOGO-E2E perform slightly worse than LINSIGHT(AUROC=0.847), one possible reason might be that LOGO has not been trained on population genomic data with conservation information to witness enough genome diversity from human and other related outgroup species. To overcome this shortcomings, LOGO-2002-C2P incorporates four evolutionary conservation features as in DeepSEA (PhastCons scores^71^, PhyloP scores^72^, and GERP++ neutral evolution^73^ and rejected substitution scores^64^) and achieves the highest performance (AUROC=0.883) (Fig. 4c) in the scenario of unrestricted negative control.

It is noted that all compared methods except DeepSEA-C2P are not specifically trained on HGMD dataset. To avoid potential over-fitting controversy and assess the generalizability of LOGO-C2P, we extract from ClinVar database^74^ to define an independent test set with 177 highly confident non-coding pathogenic SNVs (Data details in Method and Supplementary Figure 5, all splicing variants removed). LOGO-2002-C2P ranks the third (AUROC=0.927) and significantly outperforms CDTS (AUROC=0.734) but slightly worse than LINSIGHT (AUROC=0.946) and CADD(AUROC=0.966). (Fig. 4d) CDTS solely relies on 11,257 whole-genome sequences to obtain 7-mer constraint under 550-bp context of human species, whose lack of interspecies conservation leads to poorer performance to evaluate fitness consequence of inherited disease related variants. LOGO-E2E (AUROC=0.769) is only trained on 3,498 HGMD variants yet performs better than genome diversity based CDTS, which again proves end-to-end fine-tuning architecture captures some intrinsic features of non-coding genome by only using a few annotated examples. LOGO-C2P is only trained on HGMD dataset and proved to be well generalizable on ClinVar dataset. LINSIGHT is trained on human polymorphism data from 54 unrelated individuals and 3 outgroup species divergence data from aligned primates genomes conditioned on 48 genomic features, revealing the utility of incorporating genome diversity information to interpret non-coding genome. CADD is trained with more than 60 genome annotations on a much larger dataset (n=30 million) containing fixed or nearly fixed variants in human populations but is absent in human-ape ancestor as proxy-neutral variants and matched proxy-deleterious variants, which is essentially designed for binary classification of fitness consequence. The superior performance of LOGO-C2P, LINSIGHT and CADD shows that evolutionary information is likely to be powerful to identify regulatory pathogenic variants that tend to be under strong purifying selection.

We conduct another benchmark experiment to prioritize complex trait or disease associated variants. GWAS variants underlie complex diseases are generally of weaker functional impact than HGMD mutations. We construct positive test set by extracting all genome-wide significant variants (p-value<5 × 10^−8^) replicated in at least 2 independent studies from GWAS Catalog followed by retaining SNPs overlapped with ENCODE candidate cis-Regulatory Elements (ccREs)^2^ and fixation index (F_ST_) lower than 0.01 to ensure little genetic differentiation^75,76^. We ensure that all test variants have never been used in previous HGMD experiment, resulting in 2,731 positive GWAS SNPs and 704 negative controls. We bootstrap 10 times to obtain balanced held-out test set of 1,408 variants (Data details in Method and Supplementary Figure 6). All predictors show reduced performance. Compared with more deleterious HGMD mutations under significant purifying selection, common GWAS-associated variants have smaller effect size with lower evolutionary conservation, thereby plausibly more difficult to predict. LOGO-C2P-2002 is the top performer (AUROC=0.841) (Fig. 4d) across all methods We show that LOGO-C2P-2002 has the advantages of considering both chromatin effects and evolutionary constraint at base-resolution. Though LOGO-C2P is solely fine-tuned on HGMD mutations, the result proves its domain transferability from inherited diseases to common phenotypes. The second-best predictor is LOGO-919-C2P (AUROC=0.832), indicating the benefit of broad collection of chromatin features. LOGO-919-C2P outperforms DeepSEA-C2P (AUROC=0.804), which again demonstrates the edge of attention-based Transformer over CNN-based architecture. For these two independent evaluations, LOGO-C2P performs relatively better than CADD and LINSIGHT in GWAS domain than ClinVar domain, which suggests that chromatin features are more informative for complex traits while evolutionary information is more important for inherited diseases. This is consistent with the hypothesis that highly deleterious mutations of genetic diseases subject to stronger selection than complex disease loci^77^. Recent EpiMap results also emphasizes the central role of dense, rich, high-resolution epigenomic annotations to investigate regulatory circuitry of complex disease. LOGO-C2P exhibits its capability of integrating sequence context, regulatory annotation, and evolutionary constraint either explicitly or implicitly at different levels. It is noted that CDTS, which solely relies on human genetic diversity, shows poorer performance in both rare and common disease scenarios. We argue that the statistical test of 7-mer regional tolerance is not powerful enough to capture complex semantics underlying human genome sequence, even though information of more than 10,000 human genomes are incorporated^68^.

Furthermore, we explore LOGO performance on prioritizing pathogenicity of small insertion or deletion variants (Indels). We fine-tune LOGO in a similar way with LOGO-E2E using 3-mer tokenization with 1000-bp context (LOGO-E2E-Indel) on a much larger dataset from CADD Developmental release: v1.4 with 3,675,207 indels, including similar number of human-derived variants and simulations^67^. We evaluate model performance of LOGO-E2E-Indel against LINSIGHT, CADD and DeepSEA-w/o Evo (excluding evolutionary features) on independent test set, consisting of 5,869 non-coding Indels (<48bp) from ClinVar recent release (clinvar_20201003), including 5,556 positive samples (defined as pathogenic and likely pathogenic in ClinVar) and 313 negative samples (defined as benign and likely benign in ClinVar). LOGO-E2E-Indel achieves the best performance (AUROC=0.743) across all compared methods (Supplementary Figure 7). These results indicate that LOGO-E2E can effectively utilize the learned semantic representations from pre-training and shows stronger generalizability for downstream classification task than CADD, which is trained in a fully supervised manner.

## Discussion

Genome sequence contains tremendous biological information regarding the species to which it belongs. Even though a multitude of high-throughput biochemical assays have been used to characterize the sequences, the complexity nature of genome poses tremendous challenge to well interpret it. It is impractical to exhaustively perform functional annotations at every position in all conditions, and current assay design is believed to only cover the tip of the iceberg due to the limitations of existing hypothesis. A substantial gap remains between the outcomes of these experiments and a comprehensive understanding of whole genome, especially those regulatory regions. New computational approaches are in pressing needs to help interpret the underlying code. Motivated by recent huge progress in the field of NLP and CV, we propose a light language model called LOGO, utilizing ALBERT-version Transformer architecture for sequence labelling, and integrating convolution with a novel input encoding scheme for base-resolution interpretation.

Learning from raw reference genome successfully equip the model with strong adaptability across various downstream tasks by fine-tuning. No explicit annotation label is given during pre-training stage, and we have shown that the intrinsic bidirectional representations learned by the model can easily extend to sequence labeling task. In chromatin features prediction task, LOGO achieves higher accuracy than DeepSEA with significantly reduced parameters in much shorter computing time. Facing the needs of continuously growing number of functional annotations, we demonstrate supervised multitask learning faces problem of parameter explosion and tedious architecture tuning, while LOGO can efficiently extend to more abundant features with marginally increased parameters and trivial modification. Sequence-based chromatin effects prediction is informative to characterize GWAS SNPs via identifying certain regulatory function disruption. These results offer a strong justification that developing pre-trained language model can enable accurate, fast, scalable, and robust genome modelling. The community can benefit from simply and economically fine-tuning the pre-trained LOGO for specific chromatin profiles of intertest with trivial effort. By initializing model with pre-trained weights, only one additional output layer needs to be modified instead of extensive architecture tuning. We also show that fine-tuning LOGO with an explicit ref/alt token encoding strategy and convolutional operation proves powerful to prioritize functional non-coding variants associated with human disease at base-resolution.

It is noted that LOGO is only trained on human reference genome hg19. We envision that introducing genome diversity in pre-training stage can further boost representation power. This can be done by feeding LOGO with all currently identified variants across human populations and from other related outgroup species, which is expected to automatically learn evolutionary conservation and context-dependent constraint across the genome. The learned representation will in turn facilitate variant function prediction and evolutionary landscape discovery. We make an analogy between biological sequence and human language that genome possesses diversified combinations of words or phrases without compromising intrinsic grammar constraints. Overall, LOGO offers a versatile strategy to represent global and local pattern of human genome and sheds light on unearthing more value of ever-growing WGS data in the boom of national genome project.

We hypothesize that there exist many dimensions not yet captured by LOGO. There could be alternative ways to construct underlying vocabulary and define pre-training objectives with further optimized masking strategy. We already show that injecting knowledge post hoc into the model can help boost performance. On the other hand, we anticipate that large amount of existing somewhat noisy knowledgebase can be utilized to further boost the effectiveness of deep learning model. For example, sequence annotation databases^78,79^ and biological networks^80^ can be introduced systematically and structurally to guide self-supervised representation learning of genome sequence or inspire novel knowledge-guided masking strategy design. This will in turn help constructing better downstream prediction model in a more interpretable manner. In addition, LOGO can be reconfigured into a generative version, potentially be used to improve *in silico* mutagenesis efficiency and assess artificially designed sequences in the field of genome editing and synthetic biology. Integrating adversarial feedback loop of functional constraint into language model can potentially aid perturbation experiment and rational *de novo* design of new regulatory circuit^81,82^.

## Methods

### Data generation and tokenization

Coordinates in the paper refer to human genome build UCSC hg19/GRCh37. We download it from NCBI Homo sapiens reference genome with a file named GCF_000001405.25_GRCh37.p13_genomic.fna. There are five types of nucleotide, i.e., A, T, C, G and Unknown. The complementary sequence is generated using Bio.Seq (Biopython version 1.76 https://biopython.org/wiki/Seq). We divide both the forward and complementary sequence into 100-bp non-overlapping bins and obtain 60 million samples. K-mer encodes sequence into a certain length of nucleotides. For instance, a trinucleotide is a k-mer for which k=3. We define a token vocabulary to cover all possible k-mers. Token vocabulary size (V) equals to 5^k^ when using k-mer strategy. To make full use of the sequence information for rich representation, stride is designed as step-size when using k-mer to slide across a sequence. For example, for a sequence TGAATGATTTG, using 3-mer-1-stride to represent it resulting in three sets of instances, i.e., TGA ATG ATT, GAA TGA TTT and AAT GAT TTG. Then each 3-mer from each set corresponds to a token extracted from a 3-mer token vocabulary. Specifically, for a given 1,000-bp sequence, 3-mer setting results in 3 sets of input, each with length of 333 tokens; 4-mer setting results in 4 sets of input, each with length of 250 tokens; 5-mer setting results in 5 sets of input, each with length of 200 tokens; 6-mer setting results in 6 sets of input, each with length of 166 tokens. If 1,000-bp is not long enough to retrieve the last few k-mer tokens, we make up the remainder via padding forward along the reference genome.

### Architecture of the pre-training model

The pre-training model leverages the encoder part of Transformer architecture and learns representations of input sequences via multi-head self-attention mechanism. We follow the BERT notation conventions and denote the vocabulary embedding size as E, the number of encoder layers as L, and the hidden size as H. Each training instance is started with a 100-bp bin as described above and extended forward along the reference genome until reaching 1000-bp length. The model processes the genome in sequential segment of k-mer token as input, and a non-linear transformation is applied to each input token that maps input vocabulary size to a token embedding vector of length E=128. Unlike projecting the input vectors directly into the hidden space as of BERT, we leverage the optimized strategy of ALBERT, first project them into a lower dimensional embedding space of size E, and then project it to the hidden space. Token embeddings of size E are summed with position embeddings and fed into Transformer encoder network. The encoder is composed of a stack of L = 2 identical layers. Each layer has two sub-layers. The first is a multi-head self-attention layer, and the second is a position-wise fully connected feed-forward layer. A residual connection is added to each sub-layer, followed by layer normalization, leading to output of each sub-layer is LayerNorm (x + Sublayer(x)), where Sublayer(x) is the function implemented by the sub-layer itself. Each hidden sub-layer produces vector outputs with dimension of H=256. An attention function can be described as mapping a query and a set of key-value pairs to an output, where the queries (*Q*), keys (*K*), values (*V*), and output are all vectors. The output is computed as a weighted sum of the values, where the weight assigned to each value is computed by a compatibility function of the query with the corresponding key. For each self-attention layer, the input consists of queries and keys of dimension *d*_*k*_, and values of dimension *d*_*v*_. The dot products of the query with all keys, divide each by 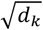, are fed into a *softmax* layer to obtain the weights on the values. The attention functions on a set of queries are computed simultaneously, packed together into a matrix *Q*. The keys and values are also packed together into matrices *K* and matrices *V*. The matrix of outputs is computed as:

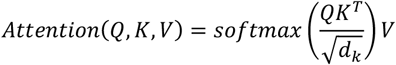

To increase the capacity of the model, the input of each hidden layer is processed by multiple attention heads, which means on each of projected versions of queries, keys and values, the attention function is performed *A* (number of heads) times in parallel. The outputs of each head are concatenated and once again projected, resulting in the final values, as depicted:

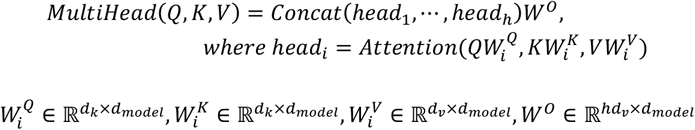

We employ *A* = 8 heads. For each of these, we use 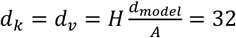. Due to the reduced dimension of each head, the total computational cost is close to that of single-head attention with full dimensionality. Following each attention layer, the fully connected feed-forward network is applied to each position separately and identically, which consists of two linear transformations with a *ReLU* activation in between.

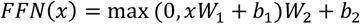

After a forward pass through L=2 layers, a final classification layer is used to project the hidden state (*d*_*model*_) to output classes of dimension equal to k-mer vocabulary size.

Motivated by ALBERT architecture, we use a factorization of token embedding parameters. By using this decomposition, the embedding parameters are reduced from *O*(*V* × *H*) to *O*(*V* × *E* + *E* × *H*). We also enforce sharing all parameters across 2 layers motivated by improved parameter efficiency of ALBERT. In original BERT model, for a given token, its input representation is constructed by summing the corresponding token, segment, and position embeddings. Position embeddings are used to capture internal relative position of each input sequence. Since we discard NSP task, we do not use segment embeddings in pre-training stage. One contribution of this work is that we demonstrate a method to incorporate prior knowledge into the language model. Knowledge layer is introduced and encoded in one-hot format. For example, if we have *M* knowledge items to label the input sequence, then a M-dimension one-hot knowledge vector is introduced and concatenated with input sequence vectors. For example, if a sequence is labelled by an annotation knowledge, all k-mers spanning from annotation start position to end position will be recorded as ‘1’ for this type of knowledge, and k-mers of other positions will be recorded as ‘0’. Knowledge embeddings are learned by the model and the dimensions are set as the same as token embeddings size. In this study, knowledge embeddings are only used in the fine-tuning stage in promoter prediction task.

### Pre-training

We define similar self-supervised loss for Masked Language Model [MLM] pre-training task as in BERT and discard Next Sentence Prediction (NSP) task. In the pre-training stage, to balance the computation burden and representation utility, we evaluate four types of k-mer (k=3,4,5,6) with 1-stride settings to tokenize the genome. For each k-mer setting of 1,000-bp sequence, k sets of input will be all used as training instances. For each set, we use similar masking strategy as in BERT. The masked token will be represented as [MASK]. We randomly masked 15% of k-mer tokens for prediction, 80% of which are replaced with [MASK], 10% is replaced by a random token from the vocabulary and another 10% remains unchanged. The original token at masked position will be predicted with cross entropy loss. The pre-training loss is the sum of the mean masked LM likelihood. We follow the BERT notation conventions and denote the vocabulary embedding size as E, the number of encoder layers as L, and the hidden size as H. In LOGO model, each k-mer of input sequence will be represented as 128-dimesion vocabulary token embedding vectors. Hidden layer embedding size (H=256) are set to be larger than input token embedding size as in ALBERT, since hidden layers are meant to learn context-dependent representations. All embeddings and model weights are expected to be learned by the model from MLM task. We use four Nvidia Tesla V100 SXM3 32G GPU to train the model. Because the number of training records exceed 180 million (3-mer: 60 million×3, 4-mer: 60 million×4, 5-mer: 60 million×5, 6-mer: 60 million×6), in order to speed up training, we convert all data to Tensorflow tfrecord and adopt Tensorflow’s *‘Multi Worker Mirrored Strategy’* strategy to support multi-machine and multi-GPU training. Parallel training technique is used on four GPUs to support large batch size. Hyperparameters are summarized as below: Layers(L)=2; Token embedding size(E)=128; Hidden dimension size(H)=256; Attention Heads(A)=8; Batch size (BSZ)=512 for each GPU, 512×4=2,048 for 4 GPUs; Steps-per-epoch=4,000; Maximum epochs=100; Sequence length=333, 250, 200, 166 tokens for 3-mer, 4-mer, 5-mer, 6-mer setting, respectively to encode 1,000-bp input sequence. We use an Adam optimizer with learning rate=0.00001. Other hyperparameters are set as default of ALBERT.

### Data for promoter prediction and fine-tuning

We use the same promoter sets from EPDnew database as of DeeReCT-PromID, which is an experimentally annotated high-quality database. We download Hs_EPDnew_006_hg19.sga from EPDnew database to construct 16,455 sequences containing human promoters and corresponding Transcription Starting Site (TSS). Promoters with (TATA+) and without TATA-box (TATA-) are also assessed independently. We make up each sequence to 10,001-bp (where +1 is a TSS position) according to hg19 reference genome as a pool of training samples. Positive sets are defined as 600-bp promoter region (from -200bp to +400bp, where +1 is a TSS position) around the known TSS. A negative control is a successive 600-bp sequence randomly sampled from all promoters’ flanking regions without overlapping (−5,000-bp to -200-bp or +400-bp to +5,000bp, where +1 is a TSS position). Since we cannot obtain original held-out test set, to make fair comparison, we use 10-fold cross-validation in fine-tuning stage and split training, validation, and test set in a ratio of 8:1:1 in each fold. The best hyper-parameters for each fold are chosen based on validation sets. We download all model weights from pre-trained LOGO to initialize the fine-tuning procedure. For each input 600-bp sequence, we tokenize them via different k-mer-1-stride settings (k=3,4,5,6) and feed them into LOGO for fine-tuning. The final hidden vector C with dimension of H corresponding to the first input token [CLS] is regarded as the aggregate representation of promoter prediction task, followed by a classification layer with sigmoid activation to obtain a score from 0 to 1 that represents the likelihood of sequence to be a promoter region. We compute a standard classification loss with C and W, i.e., log (sigmoid (CW^T^)). We use batch size of 128, set early-stop rules and train the model at most 20 epochs for each fold. We fine-tune all parameters in end-to-end manner using 1 Nvidia GeForce RTX 2060 GPU. Early stopping is used when accuracy does not improve within 3 consecutive iterations. True positive (TP), true negative (TN), false positive (FP) and false negative (FN) are defined to evaluate test set of each fold using a threshold of 0.5. We calculate three metrics to evaluate model performance, i.e., Precision, Recall and F1 score:

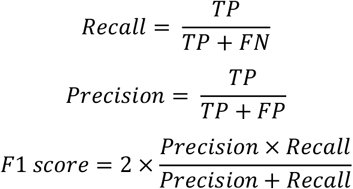

To make head-to-head evaluation, we retrain DeepReCT-PromID. We also evaluate knowledge guided fine-tuning in promoter prediction task. We download 11 annotations terms from GenBank, i.e., ‘CDS’, ‘exon’, ‘enhancer’, ‘insulator’, ‘conserved region’, ‘protein binding site’, ‘pseudogene’, ‘DNAseI hypersensitive site’, ‘nucleotide cleavage site’, ‘silencer’ and ‘gene’. Annotations of ‘promoter’ are abandoned to avoid direct label leakage. Annotation labels are constructed for each input sequence in a start-to-end spanning mode based on genome coordinate and concatenated with k-mer input. Knowledge embeddings, position embeddings and token embeddings are summed up and then fed into LOGO for end-to-end fine-tuning.10-fold cross validation is used to evaluate metric average against knowledge-naïve LOGO and DeepReCT-PromID.

### Data for promoter-enhancer interaction and fine-tuning

We use the same enhancer-promoter interaction datasets as of DeepTACT. We download all data from https://github.com/liwenran/DeepTACT, containing 65,432 permissive enhancers collected from FANTOM5 with length of 2 kb, promoter sequences defined by TSS locations from Ensembl release v75 (22) with 1kb regions surrounding each TSS, and paired enhancer-promoter interaction (EPI) labels from promoter capture Hi-C (PCHi-C) experiments in six different cell types, i.e., total B cells (tB), monocytes (Mon), fetal thymus (FoeT), total CD4+ T cells (tCD4), naive CD4+ T cells (nCD4) and total CD8+ T cells (tCD8). We use 10-fold cross validation for LOGO and DeepTACT. Due to inaccessibility of held-out test set of original DeepTACT paper, we use 10-fold cross-validation. For each fold, data augmentation technique similar with DeepTACT is applied to generate more positive training examples. We scan a certain region (say, 2 kb) surrounding the center with a 1 kb sliding window at a step size of 50 bp to obtain 20 substitutions and thus 400 pairs of positive sets. For training sets, the number of negative samples and positive samples are kept the same. For test set of each fold, we construct negative samples five times of positive ones. The detailed number of positive and negative pairs for 6 cell lines are listed in Supplementary Table 4. LOGO-EPI uses 6-mer-1-stride setting to tokenize input sequences. We add one 1D convolution operation (3 different kernel sizes) for each input sequence before fed into Transformer network. Different kernel sizes (2,3,5) of convolution layer are evaluated. Transformer output of pairing sequence are concentrated and fed into the final classification layer to obtain binary interaction label. We retrain DeepTACT to make fair comparison and use area under Precision-Recall curve (AUPRC) as performance metric. We use batch size of 128, set early-stop rules and train the model at most 40 epochs for each fold. We fine-tune all parameters in end-to-end manner using 6 Nvidia Tesla V100 SXM3 GPU for each cell-type. Early stopping is used when accuracy does not improve within 3 consecutive iterations.

### Data for chromatin feature prediction and fine-tuning

For chromatin feature prediction task, we use the same datasets as in DeepSEA and ExPecto, respectively. The genome is split into 200-bp bins as core region for each sample. To compare against DeepSEA, we pad 400-bp flanking regions on both sides to generate 1,000-bp context sequence. To compare against ExPecto with broader genomic context, we pad 900-bp on both sides to generate 2,000-bp context sequence. The 400-bp/900-bp flanking regions at the two sides provide extra contextual information to the model. Chromatin features are obtained from ENCODE and Roadmap Epigenomics projects. For each core 200-bp bin, the chromatin feature is labeled 1 if more than half of the 200-bp bin is in the peak region and 0 otherwise. Only 200-bp bins with at least one TF-binding event will be used. Forward and complementary sequence pairs share the same chromatin feature. DeepSEA contains 919 chromatin features (125 DNase I-hypersensitive sites (DHSs), 690 TF binding features for 160 different TFs and 104 histone modification mark features (HM)) and ExPecto contains 2,002 chromatin features (334 DNase I-hypersensitive sites (DHSs), 690 TF binding features and 978 histone modification mark features (HM)). We use trained DeepSEA model and ExPecto model from https://github.com/gifford-lab/deepsea-docker and https://github.com/FunctionLab/ExPecto, respectively. 919 DeepSEA chromatin features are downloaded from http://deepsea.princeton.edu/media/code/deepsea_train_bundle.v0.9.tar.gz and 2,002 ExPecto chromatin features are obtained from ExPecto author via email. Chromosome 8 and 9 are used as test chromosome. Chromosome 7 spanning the genomic coordinates 30,508,751-35,296,850 are used as validation set and not used for training. Thus, the training, validation and test sets contain 4,400,000, 8,000 and 455,024 samples, respectively. The predicted probability for each input sequence is computed as the average probability of forward and complementary sequence pairs. Performance is evaluated on test set using area under receiver operating curve (AUROC) as metric. To fine-tune on chromatin feature prediction task, we use LOGO architecture with parameter settings: L=2, E=128, H=256, A=8. The final hidden state corresponding to the first input token [CLS] is used as the aggregate sequence representation. We add a sigmoid layer after the final hidden layer of LOGO in a similar way with promoter prediction task as described above. We download model weights from pre-training stage to initialize LOGO-919 and LOGO-2002. We use 5-mer-1-stride to tokenize each input 1,000-bp or 2,000-bp sequence. Model output gives a score from 0 to 1 that represents the likelihood of sequence corresponds to each chromatin feature. Binary cross entropy loss is used to calculate loss function. We use Adam optimizer with initial learning rate of 0.0002, warmup steps=2,500, and other parameters are set as default. We use batch size of 512 with steps per epoch=4,000. We use early stopping strategy and stop training when validation loss does not further decrease within 20 epochs. We use 4 Nvidia Tesla V100 SXM3 GPU with parallelization to fine-tune LOGO-919 and LOGO-2002. We also repeat LOGO-919 fine-tuning experiment and retrain DeepSEA and ExPecto on 1 Nvidia TITAN Xp Pascal GPU to make fair comparison.

### Data processing for expanded epigenomic maps

We further expand chromatin features to more tissues and cell-lines from the latest published EpiMap results consisting of 833 high-quality epigenomes. BigWig files are downloadd from https://epigenome.wustl.edu/epimap/data/observed/. To obtain peak files, we use bigWigToBedGraph tool with default parameter settings to convert bigWig file and use MACS tool to conduct peak calling. We set parameters for MACS as *Cutoff – c*=2, *Maxgap –g*=36, *Minlen –l*=100. We obtain 2,850 ready-to-use peak files and merge with ExPecto featuresto generate 3,357 peak files after deduplicate TF-binding features. Then we use Selene library to generate training examples. At least one TF-binding event is required to extract 200-bp core regions from peak files. We randomly extract 200-bp bins from either forward or complementary chain and annotate extracted bins with 3,357 marks. We then pad 900-bp on both sides to generate 2,000-bp context sequence. Other fine-tuning settings are kept the same way as described above.

### Analysis of T2D-related GWAS variants

We download all T2D-associated SNPs from GWAS Catalog (2020-05-14 version) and obtain corresponding LD SNPs from LDlink, resulting in 156,175 SNPs (p-value ranging from 9×10^−6^ to 6×10^−447^). To make fair comparison with DeepSEA, we use the same approach to compute chromatin effects of variants. For each SNP, we extract the 1,000-bp or 2,000-bp context sequence centered on that variant based on hg19 reference genome (SNPs locates at the 500^th^ position). A pair of 1,000-bp sequence centered on both reference allele and alternative allele at the variant position are used to calculate the probabilities for each chromatin feature. Absolute differences between probability values and relative log fold changes of odds are calculated following DeepSEA pipelines. Both forward and complementary sequences are computed and averaged to obtain the predicted chromatin effects. The magnitude of the predicted chromatin effect on a chromatin feature for a SNP is computed as the product of the absolute difference between probability values and the relative log fold change of odds. We use the same protocol as in DeepSEA to obtain negative non-functional SNPs which contains 1,000,000 negative SNPs randomly chosen from 1000 Genomes Project. We calculate chromatin effects for these negative SNPs to generate the empirical background distribution and use the same E-value definition to evaluate significance of variant effects. For each chromatin feature, E-value is computed as the proportion of negative SNPs with higher predicted chromatin effect magnitude on the same chromatin feature. We use fine-tuned LOGO-919, LOGO-2000 and LOGO-3357 to calculate E-values for 919, 2,002 and 3,357 chromatin features, respectively. A variant is considered as putative functional significant if at least one chromatin feature’s E-value is equal or less than a certain threshold, i.e., 1 × 10^−5^, which might be used to infer underlying regulator disruption mechanism. Profile of Thurner islet chromatin states is downloaded from https://www.ncbi.nlm.nih.gov/pmc/articles/PMC5828664/bin/elife-31977-fig3-data5.zip. Profile of Varshney islet chromatin states is downloaded from https://theparkerlab.med.umich.edu/data/papers/doi/10.1073/pnas.1621192114/ after consultation with Dr. Narisu Narisu from Francis Collins Lab via email.

### Data processing for variant prioritization

For variants associated with inherited human diseases, we extract a dataset from Human Gene Mutation Database (version 2019-03) labelled as ‘DM’ and ‘Regulatory’ to define positive examples of regulatory variants. Within HGMD format dataset, there are some non-pathogenic alleles on complementary chain, we flip it over to forward chain to make consistent evaluation. We delete all mutations from sex chromosomes, resulting in 3,498 stringent regulatory variants. Negative variants are selected from 1000 Genomes Project SNPs. We randomly select 1 million SNPs from 1000 Genomes Project (Phase 3) and use following criteria to construct negative variants, 1) We only retain variants with allele frequency equal or greater than 0.05, 2) We delete all variants overlapped with GWAS Catalog (release 2020_05_14) reported SNPs. 3) We delete variants in RefSeq exon region and obtain 75,369 SNPs. To control allele frequency bias due to different ethnic groups, we further calculate fixation index for 75,369 SNPs. Fixation index (F_ST_) stands for a measure of population differentiation due to genetic structure and is a concrete example of Wright’s unbiased F-statistics. F_ST_ is calculated based on population size and allele frequency. According to the criteria^75,76^ that F_ST_ lower than 0.05 means little genetic differentiation and F_ST_ larger than 0.25 means high genetic differentiation. We use *vcftools* to calculate F_ST_ values for all 75,369 SNPs based on allele frequencies and sample sizes of the five super populations (AFR, EAS, EUR, SAS, and AMR) available from 1000 Genomes Project. We retain variants of F_ST_ value equal or less than 0.01 and obtain 3,690 stringent negative regulatory variants with very little population difference. We extract held-out test set of non-coding regulatory variants from ClinVar (version: ClinVarFullRelease_2020-10-03) and retain single nucleotide variants consistently annotated with ≥ two stars. We delete overlapped variants with HGMD datasets. Variants from sex chromosome and mitochondria are filtered out, which finally leads to 177 highly confident positive variants. Corresponding 177 controls are randomly chosen from above 1000G negative variants and made sure that none is used in HGMD model training. To reduce stochasticity, subsampling is performed 10 times to balance the numbers of positive and negative examples, and average performance are reported. We use GWAS SNPs associated with common human diseases or traits to construct another independent test set for variant prioritization task. We construct positive set by extracting all genome-wide significant variants (p-value<5 × 10^−8^) replicated in at least 2 independent studies from GWAS Catalog (2020-05-14 version), resulting in 7,805 SNPs. We perform further filtering via retaining SNPs overlapped with ENCODE candidate cis-Regulatory Elements (ccREs) from https://screen-v10.wenglab.org/search/?q=&uuid=0&assembly=hg19, resulting in 2,731 positive SNPs. In addition, we ensure that all sampled variants in negative control set have never been used in previous HGMD task.

### Model architecture for LOGO-E2E

To fine-tune on variant prioritization task in an end-to-end manner, we modify LOGO architecture to accommodate signed allelic information. We stack 3 layers to encode input sequence. The first layer is called ‘Ref layer’. We tokenize each 1,000-bp context sequence extracted from hg19 reference genome using 6-mer-1-stride and feed it into ‘Ref layer’ via concatenating 6 sets of 6-mer [Ref] tokens in an interlaced manner. This novel operation is introduced to encode input sequence at base-resolution without compromising representation capacity of k-mer strategy. The second layer is called ‘Alt’ layer, we use this layer to encode allelic information at certain position. Only changed position compared to ‘Ref layer’ will have input value with corresponding 6-mer [Alt] token, other positions corresponding to the context sequence are set to [Zero]. In this way, we explicitly encode the alternative allele to enforce the model to see directional alteration. The third layer is called ‘Type’ layer to encode variant type. In this paper, we do not evaluate SNVs and indels simultaneously, so we set [Type] token at corresponding position equal to 1 and other positions are set to [Zero]. Each variant with surrounding context of certain length will be encoded as a matrix input containing ‘Ref’, ‘Alt’ and ‘Type’ information, which is formatted as a ‘npz’ compressed file. 1-dimension convolutional layer is added after token embeddings and then fed into Transformer architecture. Three kernels with different sizes (2,3,5) are introduced to capture multi-scale features. Through experiments, it is found that this method reduces the weight updating frequency and makes the fluctuation range more stable during the fine-tuning process of the model. The final hidden state corresponding to learned [CLS] token embeddings is followed by a classification layer with sigmoid output of deleteriousness effect of target variant. Binary cross entropy loss is used to calculate loss function. We download LOGO-919 model weights and perform LOGO-E2E fine-tuning on HGMD training sets using 1 Nvidia TITAN Xp Pascal GPU. We use batch size of 64 and L=2, E=128, H=256, A=8. We use Adam optimizer with initial learning rate of 0.00001, and other parameters are set as default, and use early stopping strategy and stop training when validation loss no longer decreases for 3 consecutive epochs.

### Model architecture for LOGO-C2P

For LOGO-C2P, we follow similar pipelines in DeepSEA’s functional SNP prioritization architecture and firstly use previously trained LOGO-919/LOGO-2002 to generate chromatin effect features for both reference and alternative allele. We then conduct the same absolute difference and relative log fold change transformation as DeepSEA and feed these features into boosted model to train the classifier at the second stage. We assess different model performance of whether or not preserving four base-level evolutionary feature used by DeepSEA, including PhastCons scores for primates (excluding human), PhyloP scores for primates (excluding human), and GERP++ neutral evolution and rejected substitution scores. We use well-trained LOGO-919, LOGO-2002 and DeepSEA to generate chromatin features for each target variant, convert these features into DMatrix format, and train a regularized logistic regression model, using the XGBoost v0.9 implementation (https://github.com/tqchen/xgboost). It is worth mentioning that we discard z-score transformation as used in DeepSEA classifier. We argue that the tree-based approach does not require normalization as stated by original XGBoost author. The model is trained with L1 regularization parameter(alpha) =20 and L2 regularization parameter (lambda) =2,000 for iterations=1,000. Other hyperparameters are set as: Step-size shrinkage parameter(eta):0.1booster: ‘gbtree’, objective:’binary:logistic’, loss:’error’. We set early stopping rules when validation error no longer decreases for 200 epochs and preserve the best model weight. 1 Nvidia TITAN Xp Pascal GPU is used.

### Benchmarking of classifier performance on HGMD, ClinVar and GWAS variants

For HGMD regulatory variants, the performance of each model was estimated by tenfold cross-validation. For filtered 3,498 regulatory variants, we construct negative controls from 1000 Genomes Project SNPs using two schemes: random sampling (unrestricted, 3,690 negatives), and matched to positive ones within 1 kb (restricted, 3,034 negatives). We fine-tune LOGO-E2E and LOGO-C2P on HGMD dataset. We also retrain DeepSEA based classifier on HGMD dataset with or without incorporating 4 evolutionary features. CADD-precomputed scores are downloaded from http://krishna.gs.washington.edu/download/CADD/v1.3/whole_genome_SNVs.tsv.gz, FunSeq2 precomputed scores are downloaded from http://org.gersteinlab.funseq.s3-website-us-east-1.amazonaws.com/funseq2.1.2/hg19_NCscore_funseq216.tsv.bgz, GERP precomputed scores are downloaded from http://mendel.stanford.edu/SidowLab/downloads/gerp/hg19.GERP_scores.tar.gz, LINSIGHT precomputed scores are downloaded from http://genome-mirror.cshl.edu/, CDTS metrics are downloaded from http://www.hli-opendata.com/noncoding. DeepSEA functional scores are computed locally based on 919 chromatin effect predictions and 4 evolutionary information– derived scores. DeepSEA functional significance score for a variant is defined as the product of the geometric mean E-value for predicted chromatin effects and the geometric mean E-value for evolutionary conservation features. We also assess DeepSEA functional score without 4 evolutionary features. The direction of different scores for all metrics is modified to ensure lower rank represents higher probability of pathogenicity. For held-out ClinVar test set, negative controls are subsampled by bootstrapping 10 times. For held-out GWAS variants, positive samples are subsampled by bootstrapping 10 times. To compare all methods, we compute false-positive versus true-positive rates for the complete range of score thresholds. Area under the receiver operating characteristic (AUROC) is used for benchmarking.

### Benchmarking of classifier performance on CADD indels

We download the dataset from CADD Developmental release: v1.4, resulting in 3,675,207 indels. This dataset is less biased and contains much larger examples than manually curated ClinVar or HGMD. CADD is partially trained on this dataset, containing 1,837,708 *proxy-neutral* variants and 1,837,499 simulated *de novo proxy-deleterious* variants. The former sets emerge since the last human-ape ancestor and fixed in human populations, which are considered neutral (or, at most, weakly deleterious). The latter are considered free of selective pressure including both neutral and deleterious indels. We fine-tune LOGO-E2E on these CADD Indels and use human-curated dataset as held-out test set. We download known indels from NCBI/NIH ClinVar database (2020-10-03 release), which leads to 5,556 pathogenic (defined as pathogenic and likely pathogenic in ClinVar) and 313 benign indels (defined as benign and likely benign in ClinVar), respectively. Due to class imbalance, we subsample positive indels five times to construct balanced test sets and benchmark against CADD, LINSIGHT, and DeepSEA. Area under the receiver operating characteristic (AUROC) is used to benchmark different methods.

## Data availability

All datasets in this study are derived from published resources and can be generated following protocols described in Methods. Demo data is available on Github at https://github.com/melobio/LOGO.

## Code availability

LOGO is written in Python. The source code is available on Github at https://github.com/melobio/LOGO.

## Acknowledgements

We thank Zhenzhong Lan from Westlake University for discussions regarding pre-trained language model. We thank Peng Cheng Laboratory, Shenzhen, Guangdong, China to provide GPU resources to train the model. This research is supported by Guangdong Provincial Academician Workstation of BGI Synthetic Genomics (2017B090904014). Jihong Wu is supported by Program of Shanghai Academic Research Leader (20XD1401100) and Program for Outstanding Medical Academic Leader (2019LJ01).

## Author contributions

M.Y. conceived the problem and designed the study. H.M.Y and F.M. supervised the work. H.P.H and L.C.H performed deep learning computation experiments. N.Z. performed bioinformatics analysis. J.H.W provided detailed guidance on analysis of disease-related regulatory mutations. M.Y. wrote the manuscript, and other authors made modifications.

## Additional information

**Supplementary information** is available for this paper in supplementary file.

