## Supplementary Figures and Tables for "Integrating convolution and self-attention improves language model of human genome for interpreting non-coding regions at base-resolution"

##### Supplementary Figure 1

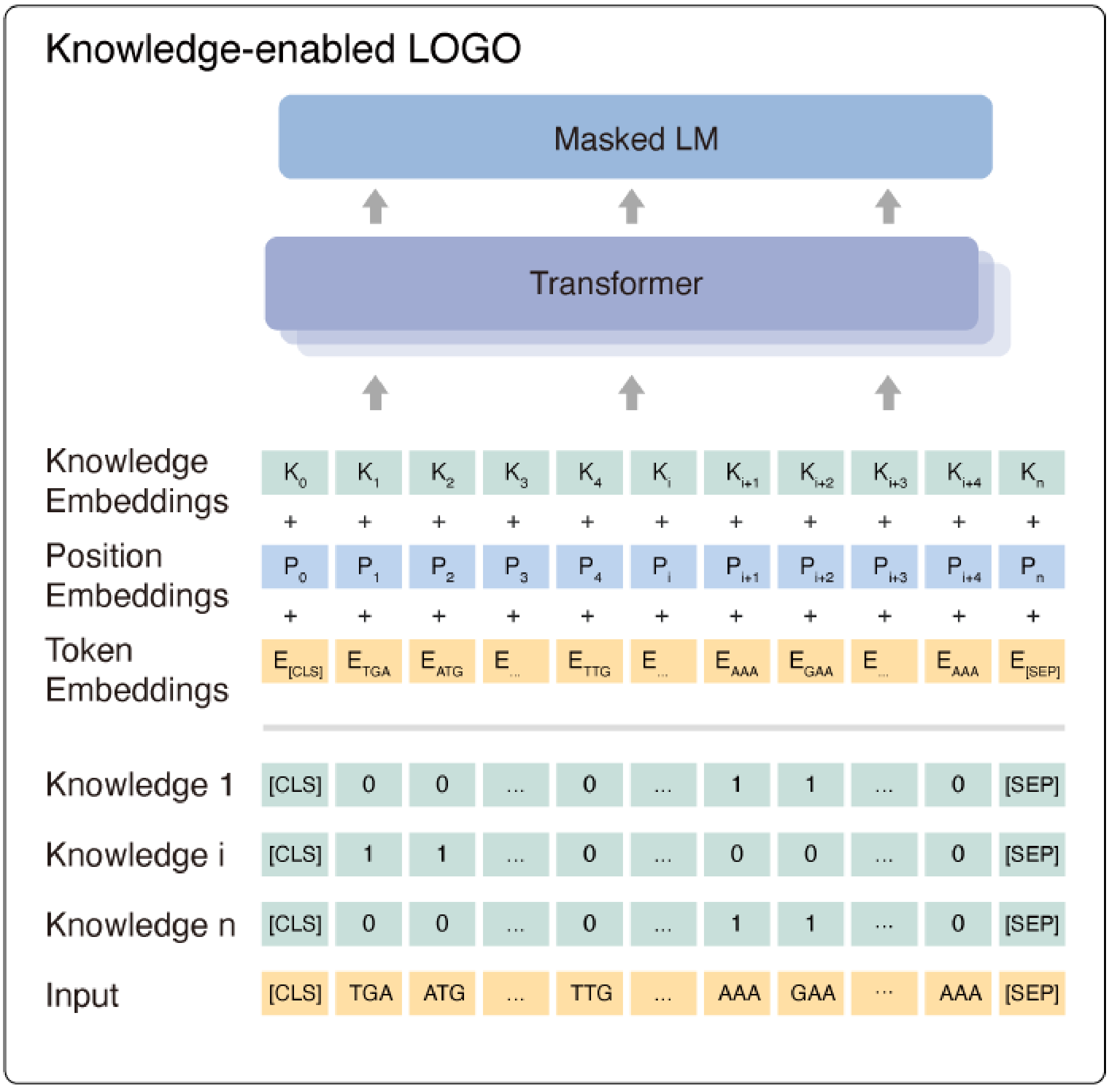


**Figure S1. Architecture of knowledge-enabled LOGO.** Knowledge layer is introduced and encoded in one-hot format. M-dimension one-hot knowledge vector represents M knowledge items to label the input sequence and concatenated with input sequence vectors. All k-mers spanning from annotation start position to end position will be recorded as '1' for this type of knowledge, and k-mers of other positions will be recorded as '0'. Knowledge embeddings are learned by LOGO and the dimensions are set as the same as token embeddings size.

**Supplementary Figure 2**

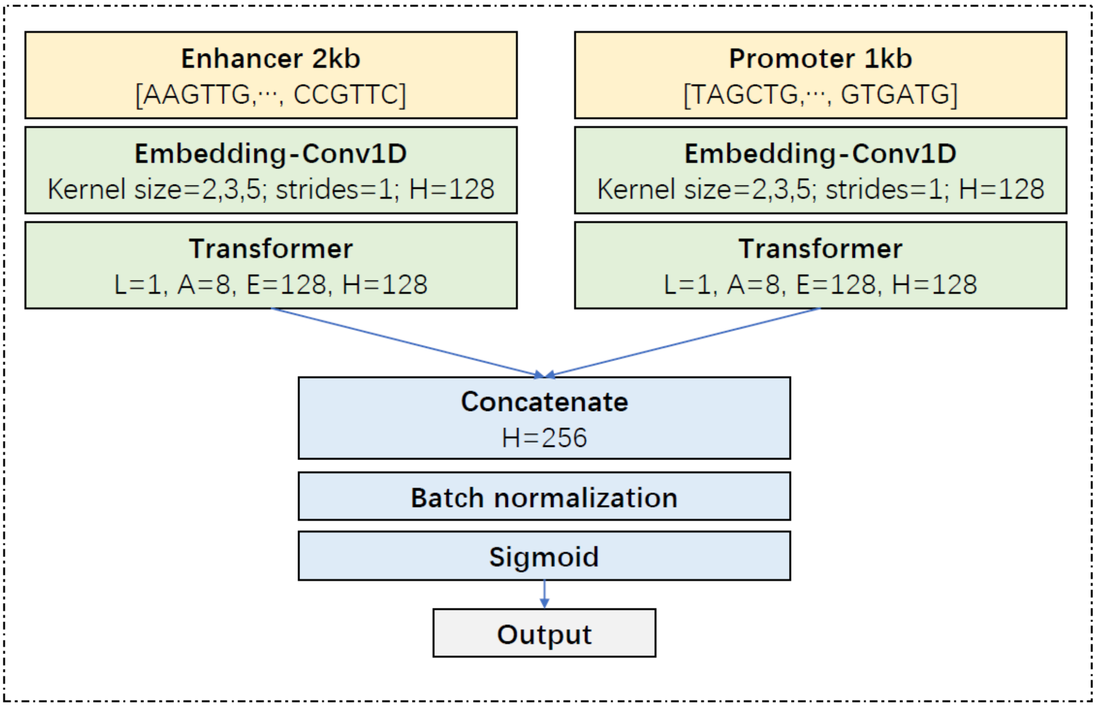


**Figure S2. Schematic diagram of LOGO-EPI's model architecture.** 1-dimension convolutional layer is added before feeding token embeddings into Transformer. Different kernel sizes (2,3,5) with 1 stride are integrated to create multi-scale features. After batch normalization processing and $Dropout = 0.25$ operation, the final interaction probability value is output. Enhancer or Promoter sequence will be truncated to fit different k-mer length. For example, when using 6-mer to tokenize 2,000-bp sequence, the enhancer sequence will be truncated into 1,980bp for round-up purpose.

**Supplementary Figure 3**

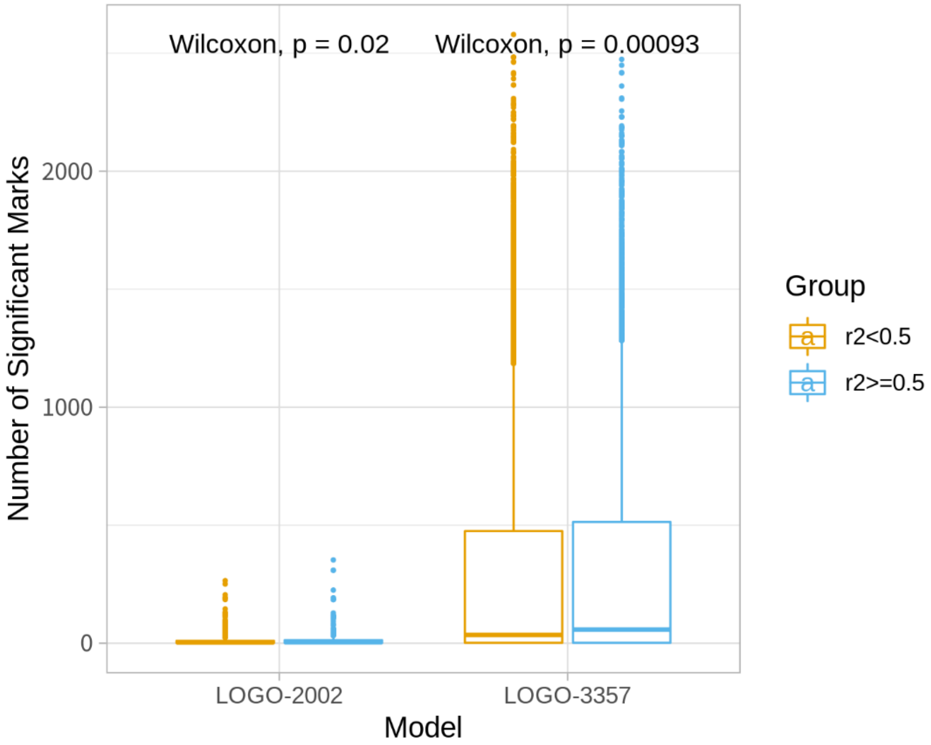


**Figure S3. The number of activated marks in 2 groups of variants with different $r^2$ values.** The difference is significant for both LOGO-2002 and LOGO-3357 (Mann Whitney U-Test).

**Supplementary Figure 4.**

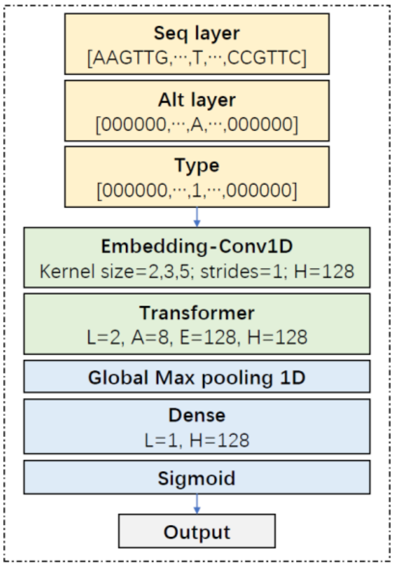


**Figure S4. Schematic diagram of LOGO-E2E's model architecture.** The first layer is called 'Ref layer'. We tokenize each 1,000-bp context sequence extracted from hg19 reference genome using 6-mer-1-stride and feed it into 'Ref layer' via concatenating 6 sets of 6-mer [Ref] tokens in an interlaced manner. The second layer is called 'Alt' layer, we use this layer to encode allelic information at certain position. Only changed position compared to 'Ref layer' will have input value with corresponding 6-mer [Alt] token, other positions corresponding to the context sequence are set to [Zero]. In this way, we explicitly encode the alternative allele to enforce the model to see directional alteration. The third layer is called 'Type' layer to encode variant type. In this paper, we do not evaluate SNVs and indels simultaneously, so we set [Type] token at corresponding position equal to [1] and other positions are set to [Zero]. Each variant with surrounding context of certain length will be encoded as a matrix input containing 'Ref', 'Alt' and 'Type' information, and then be input to 1-dimension convolutional layer. 1-dimension convolutional layer is added before feeding token embeddings into Transformer. Different kernel sizes (2,3,5) of convolution layer with 1 stride are integrated to capture multi-scale features.

**Supplementary Figure 5.**

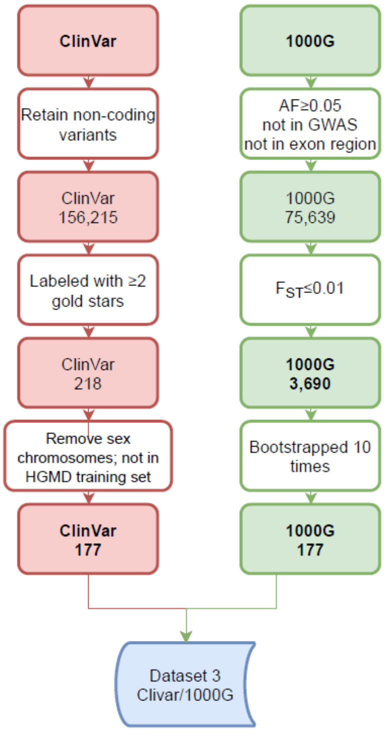


**Figure S5. Flow chart of constructing ClinVar held-out test set.** All non-coding SNVs are retained, including intergenic, intron, intronic, upstream, downstream, UTR3 and UTR5. Labeled with ≥2 gold stars, criteria provided, multiple submitters, no conflicts, practice guideline and reviewed by expert panel. Variants from sex chromosomes and duplicated with HGMD are removed, which results in 177 stringent positive variants. Balanced negative controls are subsamples from 1000G by 10 times bootstrapping.

**Supplementary Figure 6.**

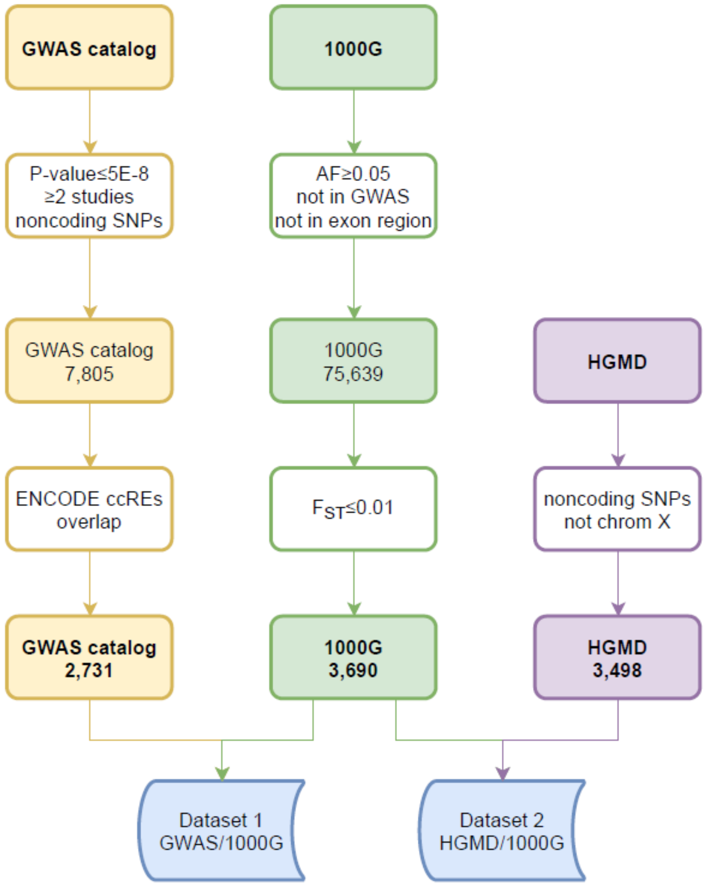


**Fig S6. Flow chart of constructing GWAS held-out test set.** Positive sets are extracted from all genome-wide significant variants (p-value$<5\times10^{-8}$) replicated in at least 2 independent studies from GWAS Catalog (2020-05-14 version), resulting in 7,805 SNPs. Further filtering is performed via retaining SNPs overlapped with ENCODE candidate cis-Regulatory Elements (ccREs), resulting in 2,731 positive SNPs. In addition, we ensure that all sampled variants in negative control set have never been used in previous HGMD task. Positive variants are subsampled 10 times by bootstrapping.

**Supplementary Figure 7.**

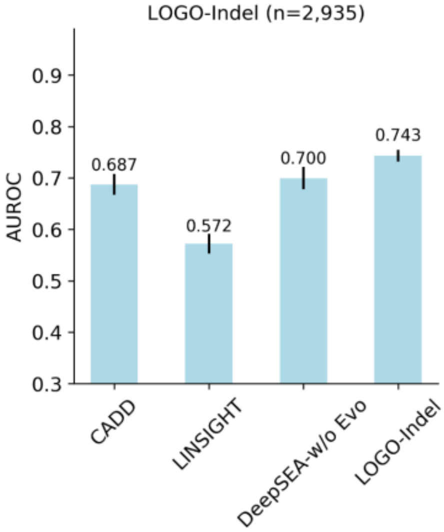


**Figure S7. Comparison of model performance by metric of AUROC on held-out ClinVar indels.** LOGO-Indel is trained on 3.2 million CADD indels and evaluated against CADD, LINSIGHT and DeepSEA-w/o Evo on held-out ClinVar indels (5,556 pathogenic indels versus 313 benign indels). Positive indels are subsampled five times to construct balanced test sets. Area under the receiver operating characteristic (AUROC) is used to benchmark different methods. DeepSEA-w/o Evo means DeepSEA derived functional significant score excluding 4 evolutionary features.

#### Supplementary Figure 8

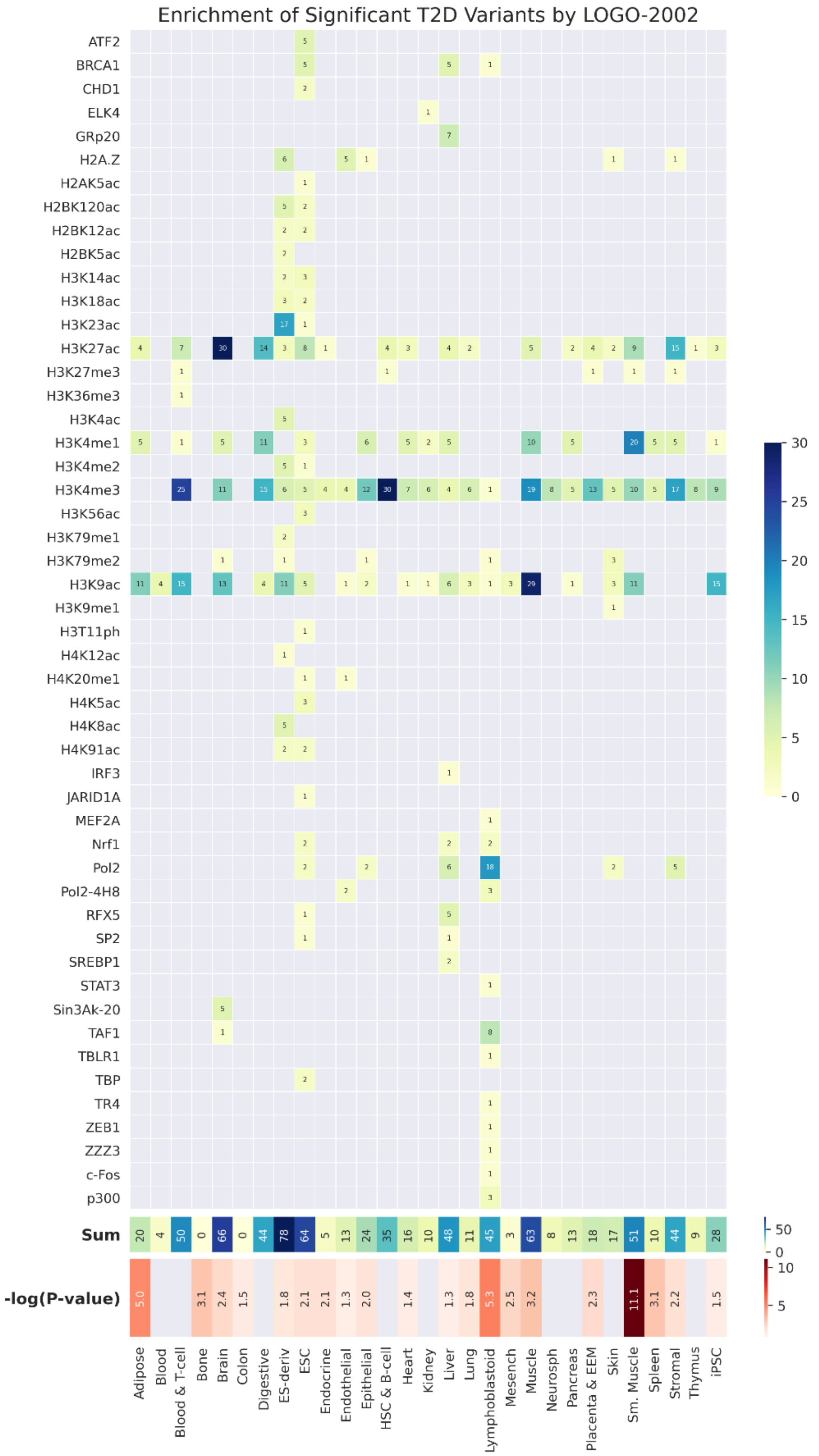


**Figure S8. Tissue enrichment analysis of significant T2D variants identified by LOGO-2002.** The number of activated chromatin marks for each type of tissue are labeled in the square. The total activation number and hypergeometric test results (-log (P-value)) for each tissue /cell type are computed and displayed on bottom panels. Only significantly enriched tissues are labeled with corresponding statistic metrics (-log (P-value)).

### Supplementary Tables

#### Supplementary Table 1

| | LOGO-3-mer | LOGO-4-mer | LOGO-5-mer | LOGO-6-mer |
|---|---|---|---|---|
| **Parameter size** | 939,786 | 1,004,286 | 1,326,786 | 2,939,286 |
| **Vocabulary size** | 125 | 625 | 3,125 | 15,625 |
| **Batch size** | 2,048 | 2,048 | 2,048 | 1,024 |
| **GPU memory (MiB)** | 17,016 | 17,016 | 17,016 | 31,172 |
| **Time per step (seconds)** | 1,870 | 2,678 | 2,954 | 2,896 |
| **Time per epoch (hours)** | 11.4 | 22.3 | 30.3 | 70.8 |

**Table S1. Hyperparameters and training configuration for LOGO pre-training.**

**Supplementary Table 2**

| Epoch | 3-mer loss | 4-mer loss | 5-mer loss | 6-mer loss | 3-mer acc | 4-mer acc | 5-mer acc | 6-mer acc |
|---|---|---|---|---|---|---|---|---|
| **1** | 0.5306 | 0.7000 | 0.8791 | 1.0310 | 0.8854 | 0.8806 | 0.8773 | 0.8800 |
| **5** | 0.4810 | 0.6258 | 0.7780 | 0.8943 | 0.8920 | 0.8872 | 0.8835 | 0.8861 |
| **10** | 0.4759 | 0.6209 | 0.7694 | 0.8890 | 0.8929 | 0.8879 | 0.8845 | 0.8865 |
| **15** | 0.4728 | 0.6191 | 0.7697 | 0.8852 | 0.8936 | 0.8881 | 0.8844 | 0.8869 |
| **20** | 0.4719 | 0.6215 | 0.7679 | 0.8867 | 0.8938 | 0.8877 | 0.8846 | 0.8866 |
| **25** | 0.4708 | 0.6161 | 0.7690 | 0.8766 | 0.8940 | 0.8885 | 0.8844 | 0.8875 |
| **30** | 0.4716 | 0.6178 | 0.7680 | 0.8830 | 0.8938 | 0.8883 | 0.8846 | 0.8869 |
| **35** | 0.4716 | 0.6210 | 0.769 | 0.8787 | 0.8938 | 0.8878 | 0.8844 | 0.8873 |
| **40** | 0.4708 | 0.6167 | 0.7673 | 0.8804 | 0.8940 | 0.8884 | 0.8847 | 0.8871 |
| **45** | 0.4676 | 0.6139 | 0.7655 | 0.8756 | 0.8947 | 0.8889 | 0.8848 | 0.8874 |
| **50** | 0.4705 | 0.6129 | 0.7632 | 0.8715 | 0.8940 | 0.8890 | 0.8850 | 0.8878 |

**Table S2. Training loss and accuracy (ACC) of pre-training LOGO using different k-mer settings.**

**Supplementary Table 3**

| Metric | Type | DeeReCT -PromID | LOGO -6-mer | LOGO -K-6-mer | LOGO -5-mer | LOGO -K-5-mer | LOGO -4-mer | LOGO -K-4-mer | LOGO -K-3-mer | LOGO -3-mer |
|---|---|---|---|---|---|---|---|---|---|---|
| Recall | TATA+ | 0.715 | 0.878 | 0.910 | 0.863 | 0.941 | 0.906 | 0.935 | 0.920 | 0.860 |
| | TATA- | 0.745 | 0.886 | 0.913 | 0.884 | 0.910 | 0.899 | 0.913 | 0.919 | 0.873 |
| | BOTH | 0.741 | 0.896 | 0.925 | 0.891 | 0.921 | 0.883 | 0.919 | 0.918 | 0.861 |
| Precision | TATA+ | 0.783 | 0.845 | 0.933 | 0.900 | 0.938 | 0.897 | 0.942 | 0.937 | 0.891 |
| | TATA- | 0.758 | 0.904 | 0.938 | 0.920 | 0.945 | 0.896 | 0.945 | 0.933 | 0.901 |
| | BOTH | 0.761 | 0.899 | 0.924 | 0.913 | 0.940 | 0.908 | 0.947 | 0.937 | 0.897 |
| F1-score | TATA+ | 0.747 | 0.861 | 0.921 | 0.875 | 0.939 | 0.902 | 0.938 | 0.928 | 0.875 |
| | TATA- | 0.751 | 0.894 | 0.925 | 0.901 | 0.927 | 0.897 | 0.929 | 0.926 | 0.886 |
| | BOTH | 0.751 | 0.897 | 0.924 | 0.901 | 0.933 | 0.895 | 0.933 | 0.927 | 0.878 |

**Table S3. Performance comparison of LOGO with different k-mer settings with or without knowledge embedding against DeeReCT-PromID.**

**Supplementary Table 4**

| Cell | FoeT | Mon | nCD4 | tB | tCD4 | tCD8 |
|---|---|---|---|---|---|---|
| **Total** | 143,617 | 174,543 | 187,973 | 189,708 | 173,867 | 175,661 |
| **Pos** | 6,676 | 8,062 | 8,712 | 9,036 | 8,282 | 8,140 |
| **Neg** | 136,941 | 166,481 | 179,261 | 180,672 | 165,585 | 167,521 |

**Table S4. Dataset details of six cell-lines for enhancer-promoter prediction task.**

**Supplementary Table 5**

| Cell | AUPRC (DeepTACT) | AUPRC (LOGO-EPI) |
|---|---|---|
| **FoeT** | 0.9372 | 0.9447 |
| **Mon** | 0.9342 | 0.9414 |
| **nCD4** | 0.9312 | 0.9457 |
| **tB** | 0.9193 | 0.9474 |
| **tCD4** | 0.9028 | 0.9475 |
| **tCD8** | 0.9364 | 0.9387 |

**Table S5. Performance details of LOGO-EPI against DeepTACT. (10-fold cross validation)**

**Supplementary Table 6**

| | LOGO-919 | LOGO-2002 | LOGO-3357 |
|---|---|---|---|
| **Pretrain/hour（Tesla V100）** | 9 | 10 | 10 |
| **Finetune/hour（Tesla V100）** | 24 | 55.3 | 117.6 |
| **Total/hour（Tesla V100）** | 33 | 65.3 | 127.6 |
| **Pretrain/hour（TITAN Xp pascal）** | 29.9 | 36.7 | 36.7 |
| **Finetune/hour（TITAN Xp pascal）** | 80 | 215.9 | 356.9 |
| **Total/hour（TITAN Xp pascal）** | 109.9 | 252.7 | 393.6 |

**Table S6. Pre-training and fine-tuning details for LOGO-919/2002/3357.**

**Supplementary Table 7**

| Model | Significant variants | Significant lead variants | Significant LD variants |
|---|---|---|---|
| LOGO-919 | 374 | 4 | 372 |
| LOGO-2002 | 729 | 7 | 727 |
| LOGO-3357 | 14764 | 71 | 14717 |

**Table S7. The number of significant T2D-related GWAS variants identified by LOGO-919/2002/3357. LD, linkage disequilibrium.**

#### Supplementary Table 8

| RS ID | Locus | Significant Marks | E-value | GWAS P-value | GWAS Odds | 1000G AF |
|---|---|---|---|---|---|---|
| rs340874 | *PROX1* | POLR2A (PNS) | 0.000006 | 1E-7 | 1.0869565 | 0.375998 |
| chr1:214159256 T-C | | | | 8E-18 | 0.0626[0.048-0.077] | |
| | | | | 3E-12 | - | |
| | | | | 6E-10 | 0.049[0.033-0.065] | |
| | | | | 1E-8 | 0.0597[0.039-0.08] | |
| | | | | 2E-22 | 1.07[1.05-1.08] | |
| rs896854 | *TP53INP1* | H3K9me3 (Blood & T-cell) | 0.000006 | 2E-6 | - | 0.515974 |
| chr8:95960511 T-C | | | | 1E-9 | - | |
| rs516946 | *ANK1* | DNase-seq (Other) | 0.000006 | 3E-22 | - | 0.804313 |
| chr8:41519248 T-C | | | | 5E-11 | - | |
| | | | | 2E-21 | - | |
| | | | | 2E-10 | 1.09[1.06-1.12] | |
| | | | | 8E-11 | 1.12[1.08-1.16] | |
| | | | | 2E-7 | 1.1[1.06-1.15] | |
| rs76549217 | *ANKH* | H3K4me2 (Endocrine) | 0.000006 | 3E-10 | 1.14[1.10-1.19] | 0.00698882 |
| chr5:14768766 C-T | | ATAC-seq (Liver) | 0.000006 | | | |
| | | DNase-seq (Kidney) | 0.000008 | | | |
| rs11583755 | *PHF13* | H3F3A (Cancer) | 0.000007 | - | - | 0.288738 |
| chr1:6672729 A-C | | H3K4me2 (ES-deriv) | 0.000009 | | | |
| rs6715901 | *TTN* | H3K27ac (ES-deriv) | 0.000007 | - | - | 0.276358 |
| chr2:179650954 A-G | | | | | | |
| rs9872347 | - | DNase-seq (Stromal) | 0.000005 | 9E-82 | - | 0.719449 |
| chr3:195831237 T-C | | BATF (Lymphoblastoid) | 0.000008 | 4E-265 | - | |
| | | BHLHE40 (Lymphoblastoid) | 0.000003 | | | |
| | | COREST (Lymphoblastoid) | 0.000010 | | | |
| | | EZH2 (Lymphoblastoid) | 0.000003 | | | |
| rs114136102 | - | CTCF (Kidney) | 0.000005 | - | - | 0.0177716 |
| chr5:36084426 T-C | | | | | | |
| rs217256 | *WNT8A* | H3K4me1 (Sm. Muscle) | 0.000008 | - | - | 0.500599 |
| chr5:137431501 T-C | | | | | | |
| rs17439448 | *SUGCT* | DNase-seq (Lung) | 0.000007 | - | - | 0.0806709 |
| chr7:40816653 T-C | | | | | | |
| rs379417 | *LINC00094* | H3K27me3 (Brain) | 0.000008 | - | - | 0.344848 |
| chr9:136890704 A-G | | IRF1 (Cancer) | 0.000008 | | | |
| | | H3K27ac (Digestive) | 0.000001 | | | |
| | | RAD21 (Cancer) | 0.000007 | | | |
| rs2482506 | *WBP1L* | EP300 (Cancer) | 0.000009 | - | - | 0.222644 |
| chr10:104563743 C-G | | H3K4me3 (Cancer) | 0.000009 | | | |
| | | H3K4me3 (Brain) | 0.000005 | | | |
| | | H3K27me3 (Digestive) | 0.000003 | | | |
| | | H2BK15ac (ES-deriv) | 0.000003 | | | |
| | | H3K4me3 (Blood T-cell) | 0.000010 | | | |
| | | H3K4me1 (Blood T-cell) | 0.000010 | | | |
| | | ATAC-seq (ATAC-seq) | 0.000006 | | | |
| rs62182438 | *HAT1* | H3K4me1 (Endocrine) | 0.000005 | - | - | 0.226438 |
| chr2:172796774 A-T | | H3K9me3 (iPSC) | 0.000002 | | | |
| | | H3K9me2 (Cancer) | 0.000008 | | | |
| | | H3K9me3 (Blood T-cell) | 0.000001 | | | |
| rs33959228 | *ZNF76* | H3K4me3 (Placenta EEM) | 0.000010 | 1E-10 | - | 0.00638978 |
| chr6:35259397 T-C | | H3K4me3 (Stromal) | 0.000009 | | | |

| | | | | | | |
|---|---|---|---|---|---|---|
| | | H3K9me3 (Muscle) | 0.000007 | | | |
| | | Max (Lymphoblastoid) | 0.000007 | | | |
| rs1327123 | - | H3K9me3 (Blood T-cell) | 0.000006 | 7E-9 | 1.04[1.03-1.05] | 0.33726 |
| chr1:184014593 C-G | | H3K36me3 (Blood T-cell) | 0.000004 | | | |
| | | DNase-seq (Blood T-cell) | 0.000008 | | | |
| | | H3K4me3 (Brain) | 0.000009 | | | |
| | | H3K27me3 (Placenta EEM) | 0.000008 | | | |
| | | H3K4me1 (Placenta EEM) | 0.000004 | | | |
| | | H3K27me3 (HSC B-cell) | 0.000005 | | | |
| | | H3K36me3 (HSC B-cell) | 0.000003 | | | |
| | | DNase-seq (HSC B-cell) | 0.000007 | | | |
| | | H3K4me3 (HSC B-cell) | 0.000002 | | | |
| | | DNase-seq (HSC B-cell) | 0.000010 | | | |
| | | H3K27ac (Epithelial) | 0.000006 | | | |
| | | H3K27me3 (Epithelial) | 0.000008 | | | |
| | | ATAC-seq (Epithelial) | 0.000010 | | | |
| | | H3K27me3 (ES-deriv) | 0.000010 | | | |
| | | H3K36me3 (Neurosph) | 0.000006 | | | |
| | | H3K9me3 (Neurosph) | 0.000009 | | | |
| | | H3K36me3 (Placenta EEM) | 0.000008 | | | |
| rs146716733 | *ARID5B* | H3K4me3 (Placenta EEM) | 0.000001 | - | - | 0.0537141 |
| chr10:63717113 T-C | | DNase-seq (ESC) | 0.000005 | | | |
| | | H3K4me3 (Stromal) | 0.000007 | | | |
| | | H3K27ac (Muscle) | 0.000008 | | | |
| | | DNase-seq (Muscle) | 0.000001 | | | |

[1] Fine-mapping type 2 diabetes loci to single-variant resolution using high-density imputation and islet-specific epigenome maps.

[2] Discovery of 318 new risk loci for type 2 diabetes and related vascular outcomes among 1.4 million participants in a multi-ancestry meta-analysis.

**Table S8. Significant SNPs (n=16) identified by LOGO-3357 overlapped with two T2D studies.**

**Supplementary Table 9**

| | Transcriptional factors (TF) | DNase I-hypersensitive sites (DHSs) | Histone modification (HM) |
|---|---|---|---|
| **LOGO-919** | 690 | 125 | 104 |
| **LOGO-2002** | 690 | 334 | 978 |
| **LOGO-3357** | 826 | 668 | 1863 |

**Table S9. Dataset details of TF/DHSs/HM for chromatin features prediction task.**
